# Antibody feedback limits the expansion of cognate memory B cells but drives the diversification of vaccine-induced antibody responses

**DOI:** 10.1101/808543

**Authors:** Hayley A. McNamara, Azza H. Idris, Henry J. Sutton, Barbara J. Flynn, Yeping Cai, Kevin Wiehe, Kirsten E. Lyke, Deepyan Chatterjee, Natasha KC, Sumana Chakravarty, B. Kim Lee Sim, Stephen L. Hoffman, Mattia Bonsignori, Robert A. Seder, Ian A. Cockburn

## Abstract

Generating sufficient antibody to block infection is a key challenge for vaccines against malaria. Here we show that antibody titres to a key target, the repeat region of the *Plasmodium falciparum* circumsporozoite protein (*Pf*CSP), plateaued after two immunizations in a clinical trial of the radiation-attenuated sporozoite vaccine. To understand the mechanisms limiting vaccine responsiveness, we developed Ig-knockin mice with elevated numbers of *Pf*CSP-binding B cells. We determined that recall responses were inhibited by antibody feedback via epitope masking of the immunodominant *Pf*CSP repeat region. Importantly, the amount of antibody that prevents boosting is below the amount of antibody required for protection. Finally, while antibody feedback limited responses to the *Pf*CSP-repeat region in vaccinated volunteers, potentially protective subdominant responses to C-terminal regions did expand with subsequent boosts. These data suggest that antibody feedback drives the diversification of immune responses and that vaccination for malaria will require the targeting of multiple antigens.

## Introduction

Induction and maintenance of humoral immunity is the mechanism of protection for most licenced vaccines against viral or bacterial infections. Most of these effective vaccines target antigenically simple infections which induce robust immune memory once the infections resolve. Even allowing for different methods to quantitate antibody titer or function, protection by these vaccines can often be mediated by relatively low amounts of specific antibodies (∼0.1-10 µg)^1^. The low amounts of antibody required for neutralization, coupled with the fact that antibody responses can have very long half-lives (10-300 years)^2^ allows our most effective vaccines to confer life-long immunity.

In contrast to current successful vaccine approaches, high antibody titers are likely to be required to protect against complex pathogens such as *Plasmodium falciparum* and HIV ^3, 4, 5^. For malaria the most advanced vaccine, RTS,S, targets the *P. falciparum* circumsporozoite protein (PfCSP) which coats the surface of the *Plasmodium* sporozoite. RTS,S given with the very potent AS01B adjuvant is administered three times at 4 week intervals and induces very high levels of antibodies (>100 μg/ml) against the immunodominant (NANP)_n_ repeat region within PfCSP; however these titers wane rapidly and this is associated with diminished protection over time ^4, 5, 6^. Anti-(NANP)_n_ repeat responses saturate after 2 immunizations and a booster at 18 months provides only a modest increase in antibody and protection^4, 5, 6, 7^.

An alternative vaccine approach has been to develop an attenuated whole parasite vaccine using radiation attenuated *P. falciparum* sporozoites (PfSPZ)^8^. This vaccine confers sterile protection in malaria-naïve individuals for ∼1 year which is thought to be mediated largely by T cells in the liver ^8, 9, 10, 11^. However, there is also evidence that PfSPZ Vaccine-induced antibodies may have some short-term protective role and utility as a correlate of protection^8, 10^.

Given the limited capacity of these current malaria vaccine approaches to induce sustained antibody mediated protection, it is critical to determine the mechanisms underlying B cell responses to *Plasmodium* sporozoites and PfCSP in particular^12, 13, 14, 15^. Analysis of antibody titer, breadth and single cell antigen specific B cell responses to RTS,S and PfSPZ vaccines in humans provides critical hypothesis generating data for developing mouse models to establish mechanisms ^8, 10, 16, 17, 18^. Here we show that following immunization with PfSPZ Vaccine in humans, B cells lose responsiveness after 2 vaccinations. To dissect the mechanism of this non-responsiveness *in vivo*, we developed Ig-knockin mice specific for PfCSP that facilitate tracking of the B cell responses to PfCSP. The data reported herein show that the lack of B cell boosting was mediated by antibody feedback by repeat-specific antibodies. However, boosting led to the emergence of subdominant epitopes and increased the diversity of the antibody response over time. This suggests that effective vaccination may depend on inducing responses to a diverse range of protective epitopes.

## Results

### Memory B cells specific for PfCSP show limited recall after 2 vaccinations

To first determine the humoral response after sequential vaccination in humans, we examined antibody responses against PfCSP in U.S. malaria-naïve adults who received 3 doses of 9×10^5^ PfSPZ Vaccine each at 8-week intervals as part of a clinical trial of this whole parasite vaccine (Figure 1A)^11^. The total anti-PfCSP antibody response increased significantly after the primary (V1) vaccination and second (V2) vaccination but did not increase following an additional boost (V3) (Figure 1B). We hypothesised that this lack of boosting could be attributed to a reduced efficiency in engaging memory B cells as part of a recall response. To investigate this, 1 week after each vaccination individual plasmablasts (PBs) from 3 randomly selected individuals were sorted and their rearranged Ig V(D)J genes amplified and cloned. Monoclonal antibodies were expressed from these rearranged Ig V(D)J sequences and screened for reactivity against PfCSP ^19, 20, 21^. After V1, only 13/153 (8.5%) PBs isolated from these 3 individuals were PfCSP-specific (Supplementary Dataset; Figure 1C). However, after V2, 45/138 (32.6%) PBs, were specific for PfCSP, many of these (29/45) used the *IGHV3-33* gene, and 18/45 belonged to one of 7 expanded clones, indicative of a robust B cell memory response at this timepoint (Figure 1C). After V3 however, only 16/112 (14.2%) PBs were PfCSP specific, suggesting that the memory B cell recall response is diminished after the first boost, consistent with the lack of increase in antibody titers at this timepoint (Figure 1C).

**Figure 1:**
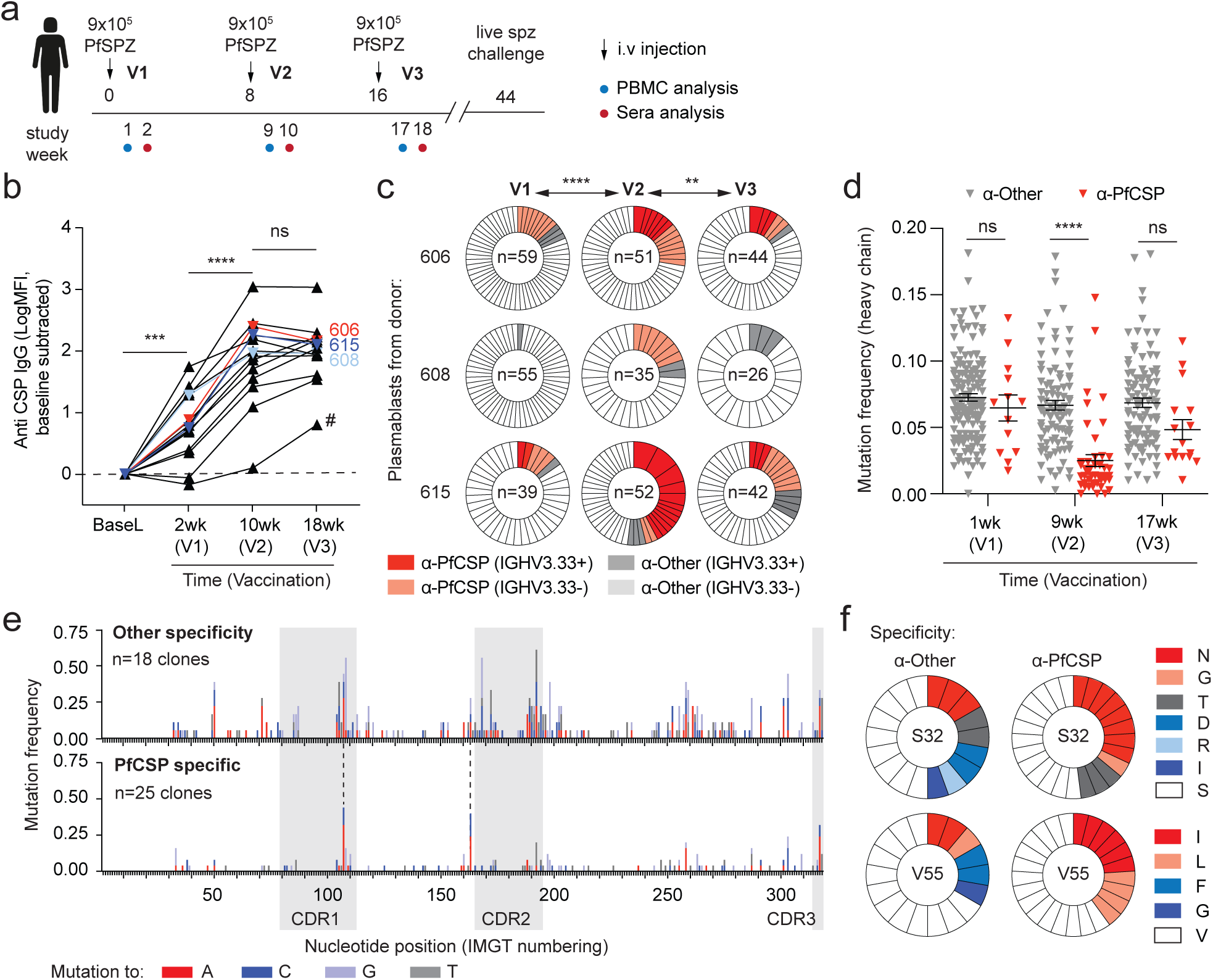
Limited memory B cell responses to PfCSP following repeated vaccination. A. Schematic of the vaccination protocol for the VRC314 clinical trial of PfSPZ with the timing of sera and PBMC collection. B. Antibody responses to whole PfCSP measured by electrochemiluminescence at the stated timepoints after each immunization, antibody responses in individuals selected for downstream PB analysis are highlighted in colour; analysis was performed by repeated measured one-way ANOVA with Tukey’s multiple comparisons test. C. Proportion of PfCSP specific PBs isolated from 3 donors after each immunization segregated by the use of the *IGHV3.33* allele; each individual wedge indicates a unique clone. Analysis was performed by chi-squared test with subject as a blocking factor. D. Mutational frequency in the heavy chain gene of non-PfCSP specific PBs (*α*-other; grey circles) and PfCSP specific PBs (*α*-PfCSP; red triangles) after each immunization, bars show mean ± s.d.; analysis by two-way ANOVA with subject as a blocking factor. E. Manhattan plots showing the location and frequency of mutations in the *IGHV3.33* genes of non-PfCSP specific PB clones and PfCSP-specific PB clones pooled from the three subjects sequenced. F. Frequency of different amino acid changes at positions 32 and 55 in PfCSP and non-PfCSP specific PB clones.

Analysis of the antibody sequences revealed that the level of somatic hypermutation in PfCSP specific antibodies was comparable to non-PfCSP binding antibodies at V1, but much lower at V2 (Figure 1D). This is consistent with the initial PB response at V1 coming from pre-existing cross-reactive memory cells as has been proposed previously^15^ while PBs recruited at V2 possibly come from memory cells that were originally primed at V1. Also in agreement with previous data^15^, we find that 8/25 PfCSP binding clones that use the *IGHV3-33* paired with a light chain formed from the *IGKV1-5/IGKJ1* in which there were no n-nucleotide insertions between the *IGKV* and *IGKJ* gene segments, resulting in a public IgL. Despite the evidence for pre-formed high affinity antibodies in multiple individuals and the low levels of SHM, many of these antibodies do appear to have undergone affinity maturation. In particular 17/25 PfCSP-specific clones that use the *IGHV3-33* gene carry mutations at either position 107 or 163 (Figure 1E; Supplementary Dataset). These mutations in the *IGHV3-33* gene commonly encode for S32N and V55I mutations at the protein level (Figure 1F). Structural analysis of a potent repeat binding antibody (mAb311) that carries both these mutations in the *IGHV3-33* heavy chain has shown that these residues are important for binding to PfCSP ^22^ further suggesting that these antibodies are under selection. Collectively, our data are consistent with specific PfCSP-binding memory cells being initially primed at V1, and then responding robustly in a secondary response to PfSPZ; however, on subsequent boosting these cells become eliminated, exhausted or limited in their expansion.

### A mouse model to investigate the PfCSP-specific B cell responses to sporozoites

To gain insight into the mechanism of the limited B cell boosting following irradiated sporozoite vaccination, we developed Ig-knockin mice – designated Igh^g2A10^ – in which the germline-reverted (unmutated) heavy chain VDJ exon (*Ighv9-3, Ighd1-3, Ighj4*) from 2A10, a murine PfCSP-neutralising antibody ^13, 23, 24^, was inserted upstream of the IgM locus (Supplementary Figure 2A). Within these mice ∼2% of the B cell repertoire is specific for the PfCSP repeat region compared with ∼0.04% of B cells within C57BL/6 controls (Figure 2A), the fact that only ∼2% of B cells in Igh^g2A10^ mice are PfCSP specific is presumably due to the fact that the inserted *Igh*^g2A10^ heavy chain remains free to pair with any endogenous light chain. Moreover, because the *Igh*^g2A10^ gene was inserted upstream of the IgM locus these antigen specific cells in naïve mice were either IgD^+^ or IgM^+^ (Figure 2A). The affinity of these cells for PfCSP was estimated by preincubation of Igh^g2A10^ splenocytes with titrated amounts of PfCSP prior to tetramer staining (Figure 2B). Based on the EC_50_ of the inhibition of tetramer staining we estimated the affinity of Igh^g2A10^ cells for PfCSP to be ∼1.33×10^−7^ M which is ∼50-fold lower that the affinity of the 2A10 mAb for PfCSP (2.7×10^−9^ M)^13^ consistent with the insertion of the unmutated germline VDJ heavy chain sequence rather than the mutated VDJ of the mature 2A10 mAb. Overall the development of total B cells was largely normal in Igh^g2A10^ mice with all spleen, bone marrow, lymph node and blood B cell populations present, though we did note a reduced number of B1a B cells and a bias towards the formation of marginal zone B cells compared to follicular B cells, perhaps because of the restricted B cell receptor (BCR) repertoire of these mice (Supplementary Figure 2B-F) ^25, 26^.

**Figure 2:**
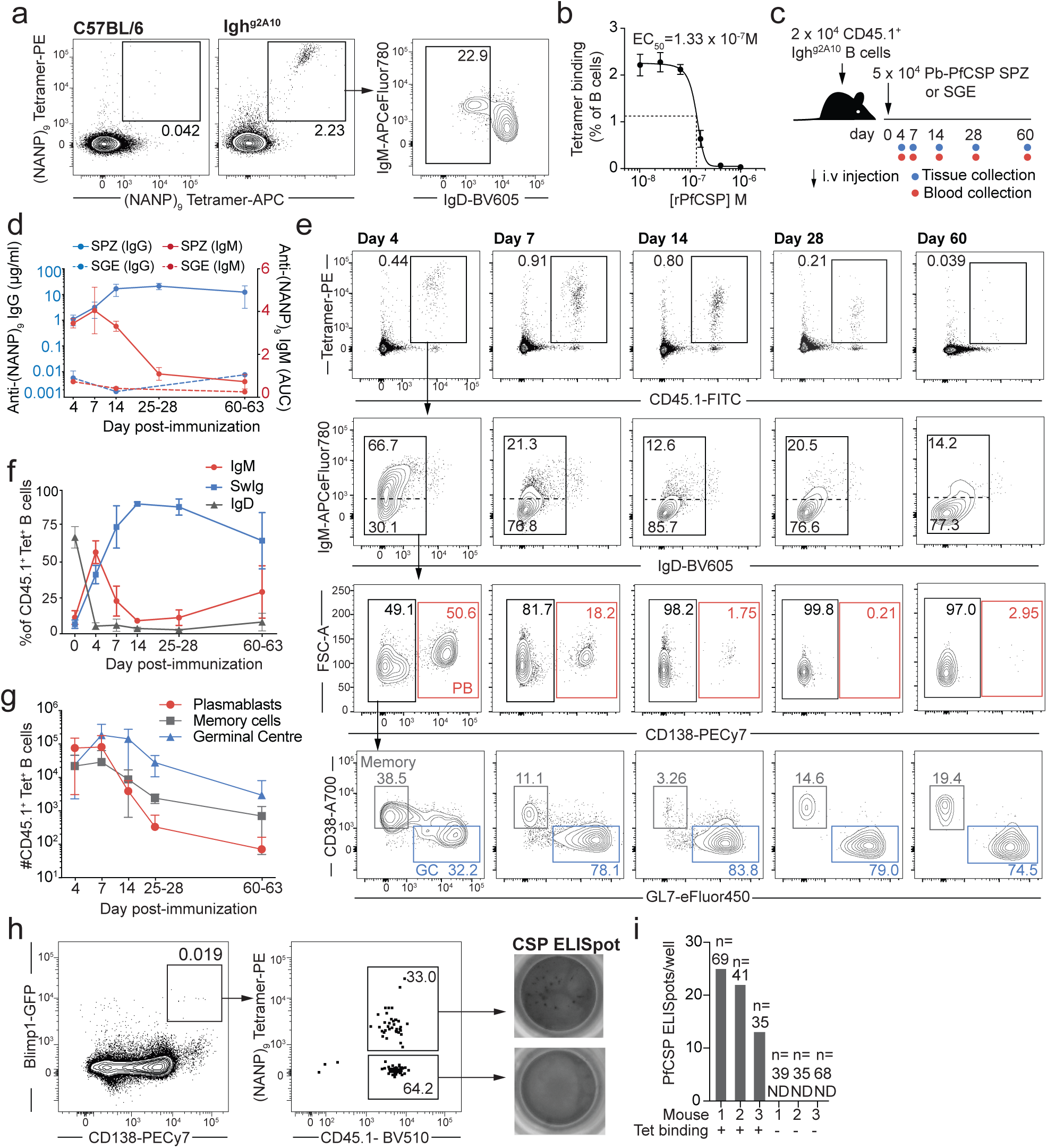
Development of Igh^g2A10^ mice to track B cell responses to PfCSP. A. Representative FACs plots (gated on B cells) showing the percentage of B cells that bind to (NANP)_9_-tetramers in C57BL/6 and Igh^g2A10^ mice, and the IgM and IgD expression on these cells. B. Titration of the concentration of PfCSP required to block tetramer staining of Igh^g2A10^ cells; Igh^g2A10^ cells were incubated with the indicated concentrations of PfCSP, prior to (NANP)_9_-Tetramer staining and flow cytometry, analysis by non-linear regression. C. Schematic of immunization experiment in which mice received Igh^g2A10^ cells and were immunized with Pb-PfCSP SPZ (SPZ) or salivary gland extract (SGE). D. Concentration of anti-(NANP)_n_ antibodies in mice immunized as in C, left axis represents IgG (expressed as μg/ml) and right axis represents IgM expressed as area under the curve (AUC), means ± s.d. shown. E. Representative flow cytometry plots showing the phenotypes of Igh^g2A10^ B cells in mice immunized with Pb-PfCSP SPZ as outlined in C. F. Summary data showing the proportions of Igh^g2A10^ cells that expressed IgD, IgM or neither (SwIg) at the indicated timepoints; means ± s.d. shown. G. Summary data showing the proportions of Igh^g2A10^ cells that are PC, GC B cells or memory B cells at the indicated timepoints; means ± s.d. shown. H. Gating strategy for the identification and sorting of Igh^g2A10^ BMPCs and representative PfCSP coated ELISpot wells probed with anti-IgG-HRP from mice immunized as in C. I. Summary data based on H. showing the number of CSP-specific Igh^g2A10^ antibody secreting cells per leg from 3 mice as identified by ELISpot, the n given above is the total number of cells in each gate that were sorted into each well.

To test whether the PfCSP-specific B cells from Igh^g2A10^ mice respond to antigen, 1 ×10^4^ congenically marked CD45.1^+^ tetramer^+^ Igh^g2A10^ B cells were adoptively transferred into naïve CD45.2^+^ C57BL/6 recipient mice, which were then vaccinated IV with radiation attenuated *P. berghei* parasites that have been engineered to express PfCSP ^27^ (Pb-PfCSP SPZ); control mice received salivary gland extract (SGE) from uninfected *Anopheles* mosquitoes as sporozoites have to be dissected from infected mosquitoes (Figure 2C). Pb-PfCSP SPZ infected mice developed early IgM responses that waned rapidly, and IgG responses that developed strongly from day 7 and reached a peak at day 25-28 of ∼20 μg/ml. No significant anti-PfCSP response was observed in mice immunized with salivary gland extract alone (Figure 2D). Following immunization with 5 × 10^4^ Pb-PfCSP SPZ, flow cytometry analysis (Supplementary Figure 2G) of the Igh^g2A10^ cells in the spleen revealed extensive expansion and class switching of the B cells after 4 days (Figure 2E and F). The early response is dominated by PBs, which subsides as germinal center (GC) and memory B cells increase, and peak, after two weeks (Figure 2E and G). Interestingly, GC B cells appear to persist for an extended period >60 days, when compared to commonly used immunisation models including HEL-SRBC and NP conjugates in which GCs resolve after ∼1 month ^28, 29^.

Because sustained antibody responses depend on the formation of a pool of long-lived bone marrow plasma cells (BMPCs) ^30, 31^, we further developed a system to facilitate the identification of these cells. Accordingly, CD45.1 Igh^g2A10^ mice were crossed to a *Blimp1*^GFP/+^ reporter mouse; Blimp1 is a key transcription factor for maintaining the plasmacell program and these mice express high levels of GFP in long-lived BMPCs^32^. In these mice, we were able to identify a population within the bone marrow, which were GFP^hi^ CD45.1^+^ and CD138^hi^ cells approximately half of which bound our (NANP)_9_-tetramers (Supplementary Figure 2H; Figure 2H). Accordingly, we sorted tetramer^+^ and tetramer^-^ CD45.1^+^ GFP^+^ cells from the bone marrow onto PfCSP coated ELISpot plates and determined that only the tetramer^+^ cells secreted PfCSP specific antibody, confirming these as antibody-secreting BMPCs (Figure 2H and I).

Finally to determine if efficient somatic hypermutation occurs in our Igh^g2A10^ cells, tetramer^+^ BMPCs were sorted and the recombined Ig V(D)J heavy and light chains sequenced using single cell RNA-seq ^33^. Single cell RNA-seq was used as we did not know the identity of the light chains in our PfCSP binding cells. *Blimp*^GFP/+^, CD45.1^+^, Igh^g2A10^ cells exclusively used the *Ighv9-3* heavy chain allele paired with the *Igkv10-94* light chain allele which is also used by the 2A10 antibody. Interestingly RNA-seq data revealed that the BMPCs were a mix of IgM^+^ and IgG^+^ cells. While ∼ 75% of IgM^+^ cells were unmutated, ∼ 90% of IgG^+^ BMPCs were somatically mutated, with all mutated cells carrying the L114F mutation in the kappa chain which has previously been shown to drive affinity maturation in the 2A10 mAb (Supplementary Figure 3). The acquisition of a low number of critical mutations indicates that affinity maturation is taking place in these B cells in a similar manner to PfCSP specific B cells in humans (Figure 1E-F). Overall the data shows that the Igh^g2A10^ knockin cells form PBs, GC B cells, memory cells and BMPCs similar to endogenous cells. Moreover, they undergo class switching and affinity maturation in a physiological manner that appears to replicate the B cell differentiation observed in PfSPZ-vaccinated humans.

### PfCSP-specific antibody and B cell responses have limited boosting in a murine model

To investigate the mechanism underlying the limited recall responses of PfCSP-specific B cells in human vaccinees following PfSPZ Vaccine, 2 ×10^4^ *Blimp*^GFP/+^, CD45.1^+^, Igh^g2A10^ B cells were transferred into congenic recipients, which were vaccinated with 5 × 10^4^ irradiated Pb-PfCSP SPZ (denoted V1, V2, V3 analogously to the clinical trials in humans) at one-month intervals between each dose (Figure 3A). Additional control groups of mice received either just one (V1) or two immunizations (V1+V2). Anti-PfCSP titers peaked after V2 at ∼70 μg/ml before declining to ∼40 μg/ml and, similar to vaccinated humans, this response was not enhanced by a further boost (Figure 3B). We further analysed the PB, GC and memory cell responses at the cellular level by flow cytometry (Figure 3C and D). We detected a transient expansion of the PB (CD45.1^+^, CD138^hi^, GFP^lo^) and memory (CD45.1^+^ CD38^+^) response after V2 but not V3 (Figure 3E and F); however, there was no change in the ongoing GC response (Figure 3G). Boosting was associated with a small increase in BMPCs after V2 that was marginally significant, however a third boost did not enhance numbers further (Figure 3H). Taken together, these data are consistent with the observations in humans (Figure 1B and C) that memory cells can be recalled into the response at V2, but not at V3 limiting the titer of neutralizing antibody that can be achieved.

**Figure 3:**
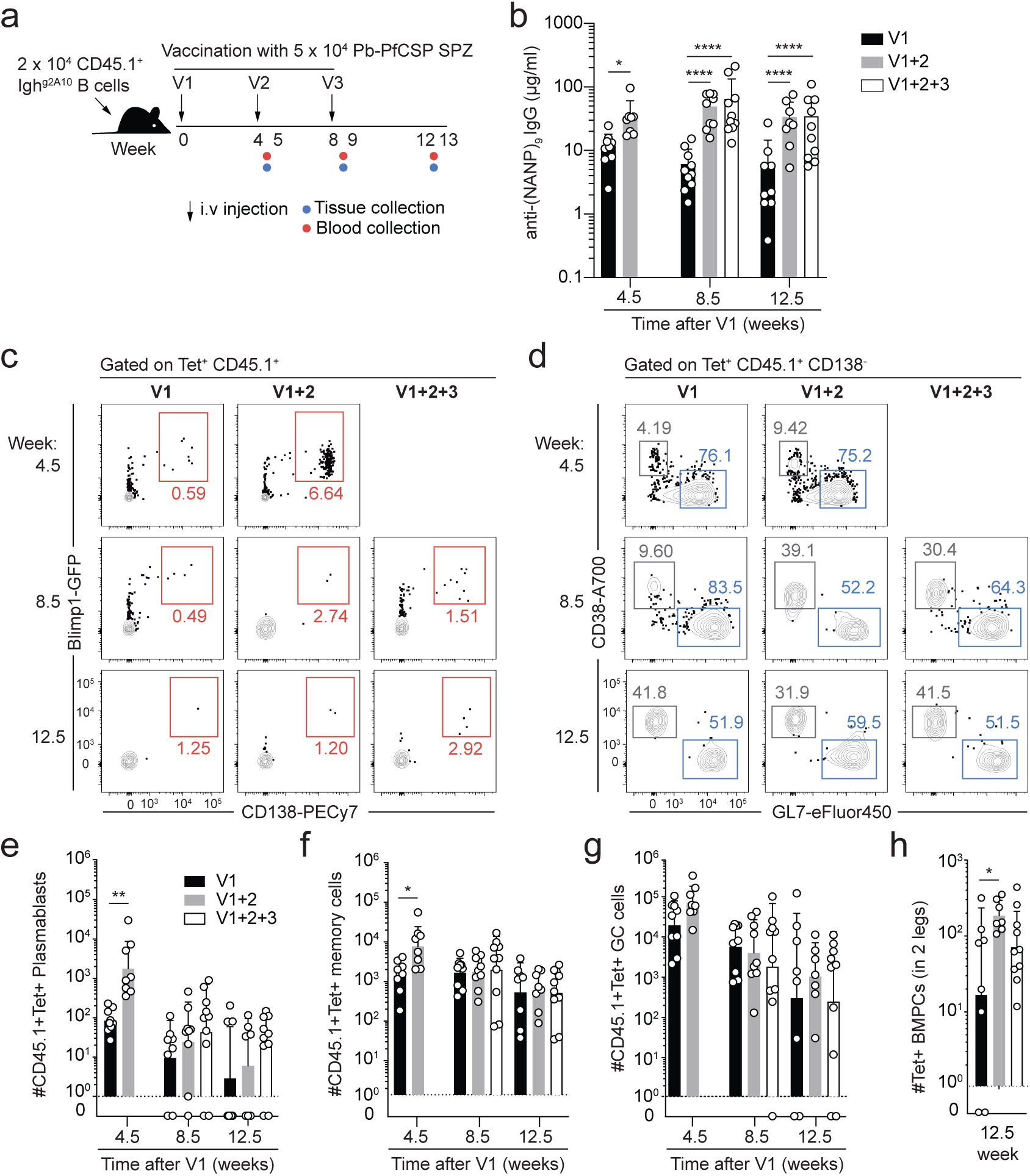
Limited memory B cell recall responses to PfCSP following repeated vaccination in mice. A. Immunization schedule for the experiment: mice received 2 × 10^4^ congenically marked Igh^g2A10^ Blimp^gfp/+^ cells and were immunized 1, 2 or 3 times with 5 × 10^4^ Pb-PfCSP SPZ at 4 week intervals. 5 days after each boost and 33 days after the final boost blood, splenocytes and bone marrow was collected from the mice for analysis by ELISA and flow cytometry. B. Concentrations of anti-(NANP)_n_ IgG in the sera at the indicated timepoints. C. Representative flow cytometry plots for the identification of Igh^g2A10^ PBs and plasma cells in the spleen. D. Representative flow cytometry plots for the identification of Igh^g2A10^ GC B cells and memory cells in the spleen. Summary data for the analysis of spleen PBs (E), spleen memory B cells (F), spleen GC B cells (G) and BMPCs (H) at the indicated timepoints; data are pooled from 3 replicate experiments, analysis was by 2-way ANOVA including the experiment as a blocking factor, bars represent means ± s.d..

### Sporozoite-induced memory cells are functional

The next series of studies determined why the recall response was so rapidly diminished. One hypothesis is that Pb-PfCSP SPZ immunization induces non-responsive memory B cells. To directly test this, memory cell populations were generated by transferring Igh^g2A10^ B cells to congenic mice and immunizing with either Pb-PfCSP SPZ or recombinant PfCSP (rPfCSP) in alum as a proxy for a recombinant protein immunization. After >2 months, switched memory B cells which expressed PD-L2 and CD80, markers of functional memory B cells were detected ^34^, suggesting normal differentiation (Supplementary Figure 4A and B). Negative selection was then used to enrich antigen experienced memory B cells from immunized mice and remove contaminating plasma cells and GC B cells (Supplementary Figure 4C). The functional capacity of the memory B cells was assessed by adoptive transfer into naïve mice and re-immunization with 5 × 10^4^ Pb-PfCSP SPZ (Figure 4A). Immunized mice that had received memory cells had significantly higher titers of PfCSP-specific IgG, but not IgM, than immunized mice that did not receive cells (Figure 4B), indicating that the transferred memory cells could differentiate into antibody secreting cells upon recall. At the cellular level, memory cells from Pb-PfCSP SPZ immunized mice expanded ∼50 fold in 5 days, and differentiated into both CD138^+^ PBs and GL7^+^ GC B cell precursors (Figure 4C-E). This level of expansion was only slightly lower that that seen among memory B cells primed with rPfCSP in Alum. Overall, our experiments indicate that irradiated sporozoite immunization induces memory cells that appear capable of mounting robust recall responses.

**Figure 4:**
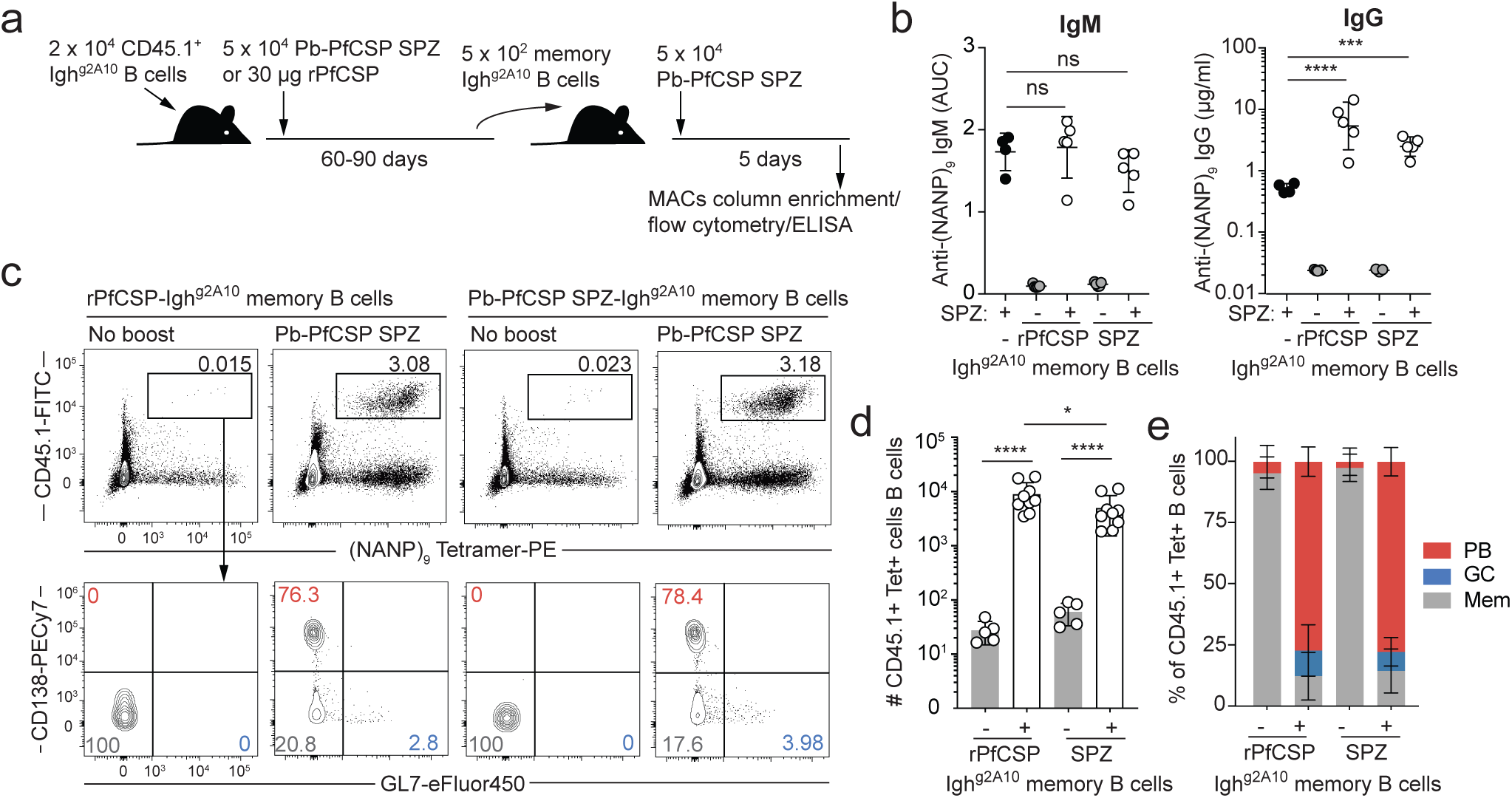
Memory B cells induced by sporozoite immunization are able to mount recall responses. A. Schematic of the experiment showing the protocol for the generation and transfer of Igh^g2A10^ memory B cells to naïve recipient mice and subsequent boosting with Pb-PfCSP SPZ. B. Levels (NANP)_n_-specific IgM, and IgG in mice that received Igh^g2A10^ memory B cells 5 days after boosting with 5 × 10^4^ Pb-PfCSP SPZ compared to naïve mice that did not receive memory cells, but were immunized concurrently with 5 × 10^4^ Pb-PfCSP SPZ; data from a single experiment, bars show mean ± s.d., analysis by one-way ANOVA with Tukey’s multiple comparisons test. C. Representative flow cytometry plots of Igh^g2A10^ memory cells recovered by magnetic bead purification from recipient mice immunized as in A. gated on CD19^+^ or CD138^+^ B cells and PBs. D. Quantification of the numbers of recovered cells in each group; bars show mean ± s.d., data pooled from 2 experiments with analysis by two-way ANOVA with experiment as a blocking factor. E. Proportions of memory cells from (D) that had differentiated into PBs, GC B cells or retained a memory phenotype (Mem); means ± s.d. shown.

### PfCSP specific memory B cells are inhibited by antibody feedback

Given that the PfCSP-specific memory B cells were functional in naïve mice, we next investigated whether they were regulated by other components of the ongoing immune response. Accordingly, memory B cells were adoptively transferred 1 month after immunization into immune-matched mice that had received a Pb-PfCSP SPZ immunization (Figure 5A). In immune-matched mice, the expansion of transferred memory B cells was significantly limited, with only a small number of cells differentiating into PBs (Figure 5B-C). This regulation did not appear to be mediated by the cellular response as Pb-PfCSP SPZ pre-immune MD4 mice, which have an Ig specific for hen egg lysozyme and should not produce PfCSP-specific antibody^35^ but have normal T cell responses, failed to show the same inhibition as the pre-immune C57BL/6 mice. Further, passive transfer of immune sera was sufficient to severely limit the memory B cell response (Figure 5B-C). Overall the inhibition of memory B cell expansion appeared to correlate with the level of anti-(NANP)_n_ antibodies (Figure 5D). If antibody feedback was the mechanism limiting memory responses in human PfSPZ vaccinees, there should be an inverse relationship between the amount of antibody prior to a booster immunization and the subsequent change in the antibody titer. In agreement with this, there was a strong inverse correlation between post V2 antibody titres and V3 boosting (r^2^ = 0.51; p = 0.0041) in vaccinated humans, indicative of strong antibody feedback regulation limiting the third immunization (Figure 5E). Conversely, no such relationship was observed at the earlier V2 immunization (r^2^ = 0.12; p = 0.22), indicating that antibody feedback was not strongly regulating B cell responses at this earlier timepoint.

**Figure 5:**
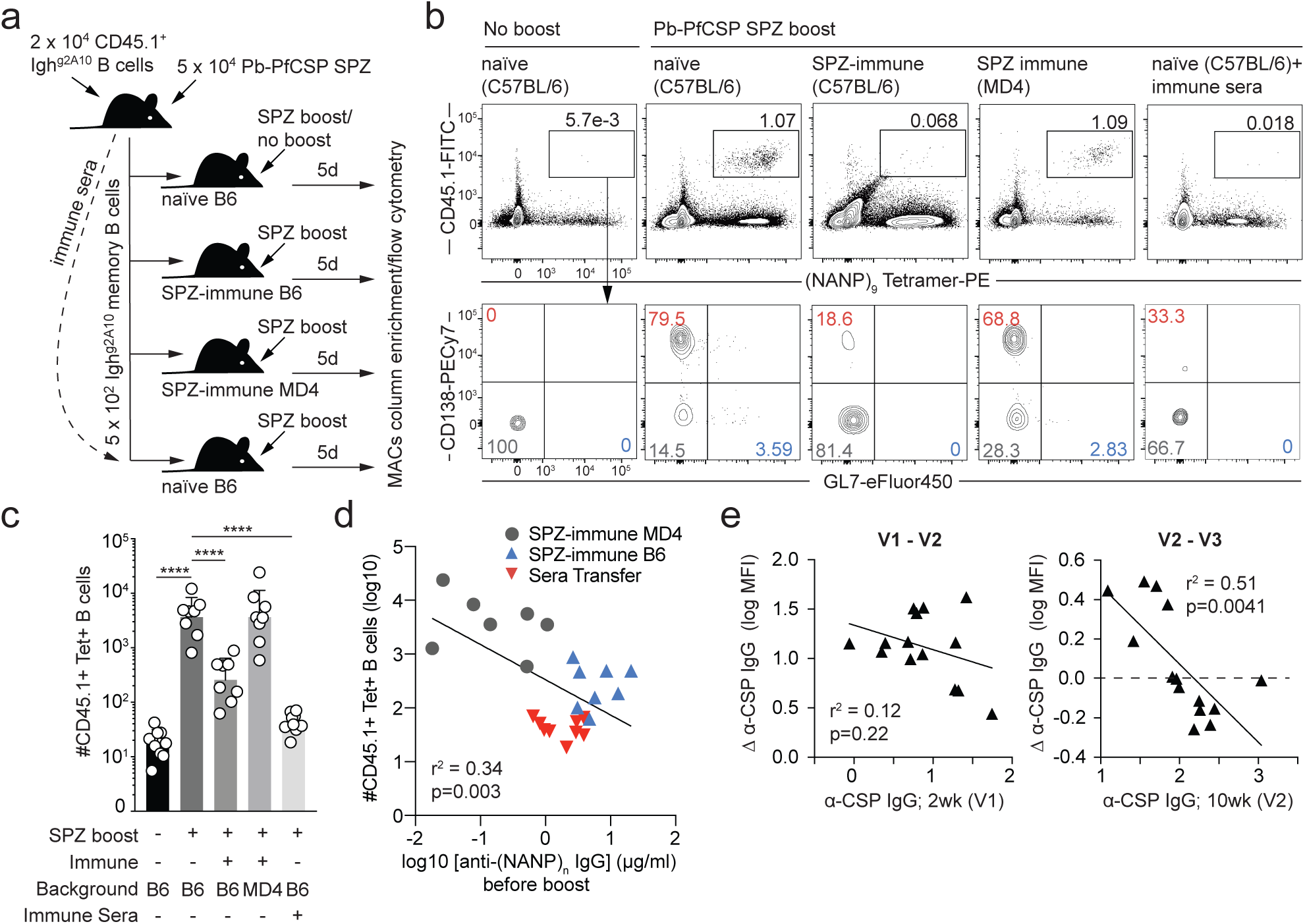
Antibody feedback regulates memory B cell responses in mice and humans. A. Schematic of the experiment showing the generation of and transfer of memory cells and sera to different groups of naive and immune recipients prior to boosting with 5 × 10^4^ Pb-PfCSP SPZ. B. Representative flow cytometry plots of Igh^g2A10^ memory cells recovered by magnetic bead purification from recipient mice immunized as in A. gated on CD19^+^ or CD138^+^ B cells and PBs. C. Quantification of the numbers of recovered cells in each group; data pooled from 2 experiments, bars show mean ± s.d., with analysis by one-way ANOVA using Tukey’s multiple comparisons test, with experiment as a blocking factor, only significant comparisons with the positive control group (Pb-PfCSP boosted naïve C57BL/6 recipients) are shown. D. Correlation of the sera titers of anti-(NANP)_n_ antibodies prior to boosting in the different groups of mice in A. with the subsequent size of the expansion of the CD45.1^+^ Tetramer^+^ B cell population after boosting; data pooled from 2 experiments, analysis by linear regression. E. Correlation of the response (change in anti-PfCSP antibody level) to V2 and V3 boosts with the titer of antibodies prior to the corresponding boost among the PfSPZ vaccinated individuals described in Figure 1A; analysis by linear regression.

We further predicted, that if antibody feedback regulates recall responses, then delaying boosting until antibody titers have declined would enhance recall responses. Accordingly, in two groups of mice we delayed the first boost until 6 months after the initial priming immunization (Supplementary Figure 5A). In these mice titres peaked at ∼100 μg/ml (Supplementary Figure 5B), which was higher than the peak in mice that received a boost at 1 month (∼70 μg/ml; Figure 3B). The magnitude of the secondary PB response was not obviously larger in these mice than in mice that received a boost one month after the prime (Supplementary Figure 5C-D), but the delayed mice exhibited secondary GCs and so the B cells may have enhanced SHM and affinity maturation resulting in higher titers of protective antibody (Supplementary Figure 5E-G).

### Sub-protective levels of anti-repeat antibodies block recall responses by memory B cells

For vaccination a critical metric is the protective threshold of antibody required for protection. If this protective threshold is above the amount of antibody required to inhibit memory B cell responses it will be more difficult to achieve sustained protective titers by vaccination. Accordingly, we determined the protective threshold for the 2A10 antibody by passively transferring different amounts of antibody to mice and subsequently challenging them via bites of Pb-PfCSP infected mosquitoes and following mice for 14 days or until they became infected (>0.1% parasitaemia). A dose of 300 µg provided complete protection in 7/10 mice, but a dose of 100 μg conferring only partial protection (3/10 mice protected; Figure 6A). Passive transfer of 100μg antibody resulted in a serum concentration of ∼70 μg/ml (Figure 6B) which is approximately double the amount achieved by vaccination in our mice (Figure 3B) and similar to the level of antibody required for 50% protection following RTS,S vaccination ^4^. We next determined the dose of antibody required to inhibit recall responses in mice by transferring memory B cells to mice and subsequently boosting with Pb-PfCSP SPZ in the presence or absence of antibody. 2A10 antibody inhibited B cell responses at all concentrations tested (Figure 6C), with the lowest concentration of 33 μg permitting only minimal differentiation of CSP specific B cells (Figure 6D). These data are consistent with the fact that there was no response at V3 in the mice at a timepoint when the serum titer of antibody was ∼70 μg/ml (Figure 3C).

**Figure 6:**
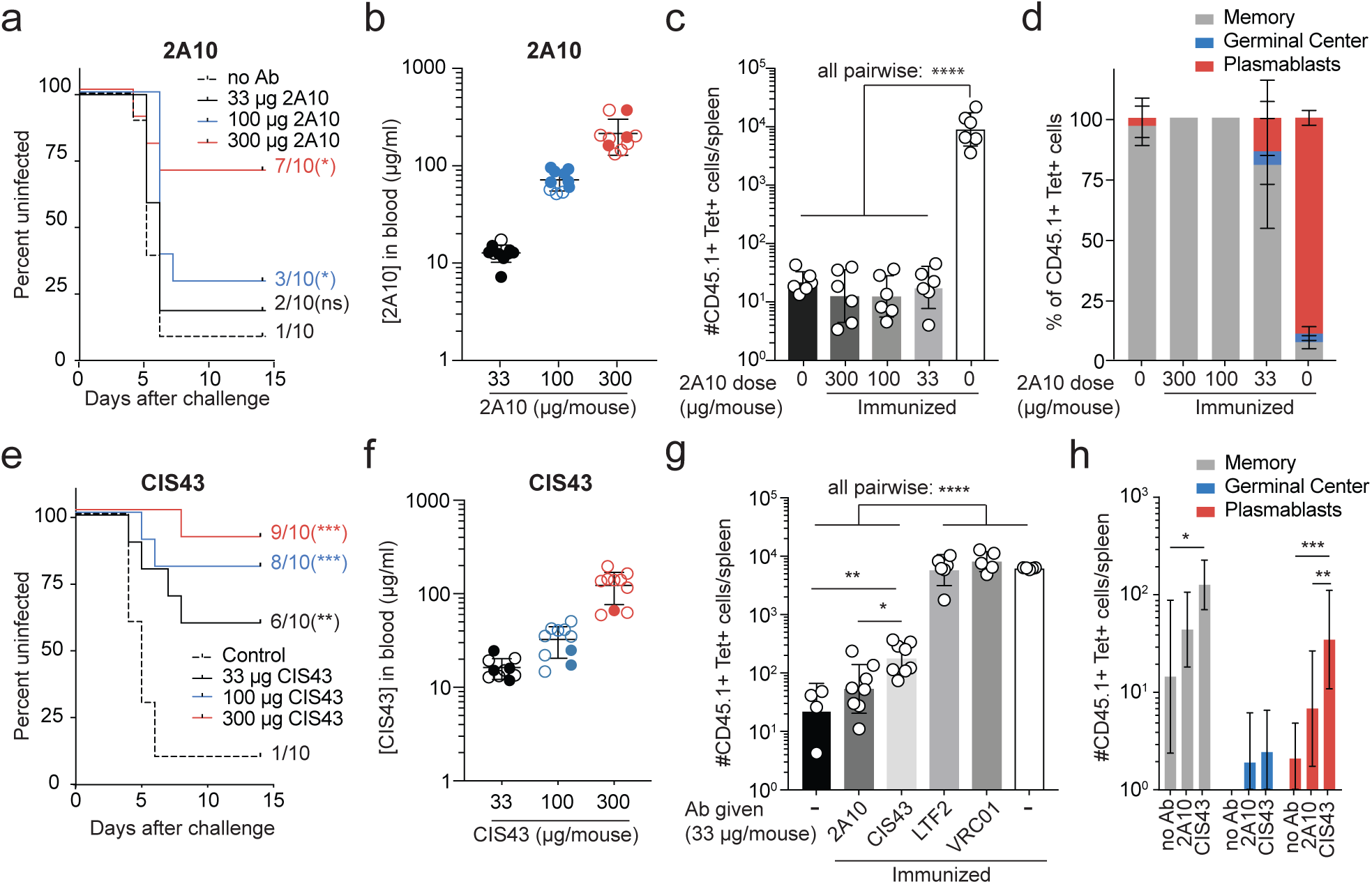
Sub-protective levels of antibody potently inhibit memory B cell responses. A. Survival plots showing the proportion of uninfected mice after IV transfer of the specified amounts of 2A10 antibody and feeding by 7 infected mosquitoes; data pooled from 2 experiments, analysis by Log-rank (Mantel-Cox) test showing pairwise comparisons with the no antibody group. B. Concentration of 2A10 antibody in the blood 2 hours post-transfer via ELISA; hollow circles indicate protected mice, means ± s.d. shown. C. Expansion of Igh^g2A10^ memory B cells (generated using Pb-PfCSP SPZ immunization and boosted as in figure 4A) in recipient mice that received the specified doses of 2A10; data pooled from 2 independent experiments and analysed by one-way ANOVA using Tukey’s multiple comparisons test with experiment as a blocking factor. D. Proportions of Igh^g2A10^ memory cells from (C) that had differentiated into PBs, GC B cells, or retained a memory phenotype. E. Survival plots showing the proportion of uninfected mice after IV transfer of the specified amounts of CIS43 antibody and feeding by 7 infected mosquitoes; data pooled from 2 experiments, analysis by Log-rank (Mantel-Cox) test showing pairwise comparisons with the no antibody group. F. Concentration of CIS43 antibody in the blood 2 hours post-transfer via ELISA; hollow circles indicate protected mice, means ± s.d. shown. G. Expansion of memory B cells (generated using Pb-PfCSP immunization and boosted as in figure 4A) in recipient mice that received 33 μg of the specified anti-CSP (2A10 and CIS43) antibodies, and antibodies of irrelevant specificity (LTF2 and VRC01); data pooled from 2 independent experiments and analysed by one-way ANOVA using Tukey’s multiple comparisons test with experiment as a blocking factor, means ± s.d. shown. H. Numbers of recovered PB, GC and memory CD45.1+ Tetramer+ cells from the no antibody/no immunization, 2A10 and CIS43 groups in G; means ± s.d. shown; analysis by two-way ANOVA with experiment as a blocking factor.

To determine whether this effect is generalizable to other anti-PfCSP antibodies the ability of the potent human neutralizing antibody CIS43 to inhibit the expansion of Ighg2A10 was investigated. CIS43 has dual binding activity with affinity for both the (NANP)_n_ repeat and the junction between R1 and the repeat^20^. In broad agreement with our previous work we found that concentrations of ∼15 μg/ml blocked around 50% of infections, significantly better than 2A10 (Figure 6E-F). These serum concentrations also potently inhibited Igh^g2A10^ memory B cell expansion, upon boosting, compared to mice that received no antibody, or control antibodies with irrelevant specificities but not to the same degree as 2A10 (Figure 6G). Moreover, some of the memory B cells that did expand were able to differentiation into PBs in the presence of semi-protective levels of CIS43 (Figure 6H).

### Antibody feedback occurs via epitope masking permitting the expansion of subdominant responses

The final experiments were aimed at determining the mechanism of feedback inhibition by antibody. In particular, whether antibody inhibition regulated PfSPZ specific B cell responses generally or if the inhibition was antigen specific. General mechanisms of antibody feedback would include action via inhibitory Fc receptors^36^, or clearance of parasite before parasite antigens could enter presentation pathways^37^. Alternatively, if the antibodies were acting via epitope masking the feedback would be epitope specific such that antibodies specific for other regions of the CSP molecule would not inhibit responses by our (NANP)_n_ -repeat specific cells (Figure 7A).

**Figure 7:**
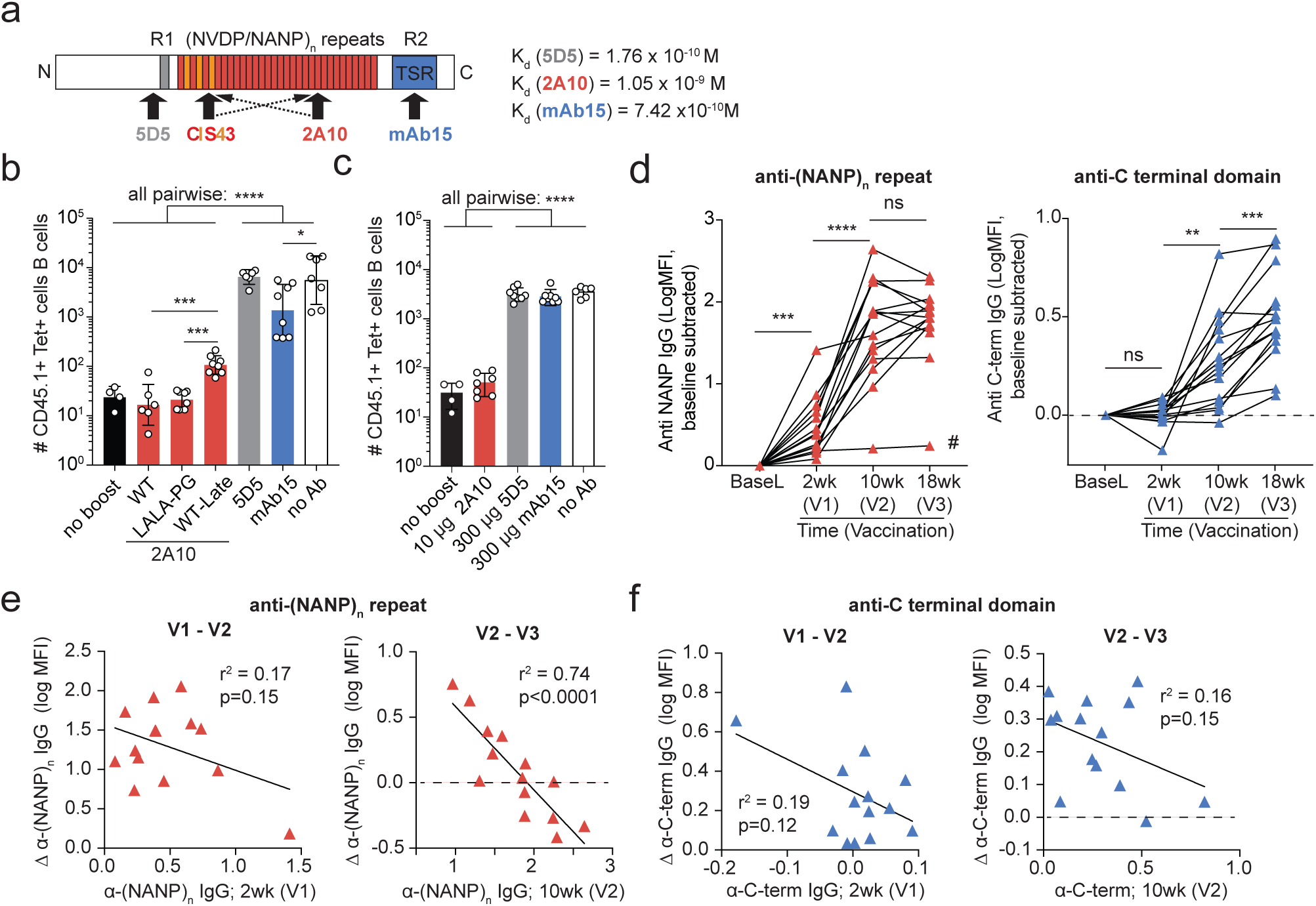
Antibody feedback occurs via epitope masking and allows the diversification of the antibody response. A. Schematic of the CSP molecule showing the binding sites and dissociation constants of the different antibodies used in this study. B. Expansion of memory B cells in the presence of antibodies (30 μg/mouse) targeting non-repeat regions of CSP (5D5 and mAb15), anti-repeat Fc-dead 2A10 (2A10-LALA-PG) and 2A10 transferred 4 hours post sporozoite delivery; memory cells were generated via Pb-PfCSP SPZ immunization and transferred as in figure 4A and expansion measured 5 days post boosting with 5 × 10^4^ Pb-PfCSP SPZ. C. Expansion of memory B cells in the presence of anti-PfCSP antibodies at different concentrations; memory cells were generated via Pb-PfCSP SPZ immunization and transferred as in figure 4A and expansion measured 5 days post boosting with 5 × 10^4^ Pb-PfCSP SPZ. D. Antibody responses specific for the (NANP)_n_-repeat and C-terminal domain of CSP in PfSPZ vaccinated subjects (described in Figure 1A) analysis was performed by repeated measures one-way ANOVA with Tukey’s multiple comparisons test; # indicates one individual who did not respond and was excluded from subsequent analysis. F. Correlation of the response (change in anti-(NANP)_n_ antibody level) to V2 and V3 boosts with the titer of antibodies prior to the corresponding boost; analysis by linear regression. G. Correlation of the response (change in anti-terminal antibody level) to V2 and V3 boosts with the titer of antibodies prior to the corresponding boost; analysis by linear regression.

To determine if inhibitory Fc receptors were important, 2A10 LALA-PG mutant antibodies carrying Leucine-to-Alanine mutations in positions 234 and 235, and a Proline-to-Glycine substitution at position 329 in the Fc portion of the antibody were made that limit binding with Fc-gammaRIIB receptors^38^. These antibodies, however, inhibited the expansion of memory cells similarly to the unmutated antibodies (Figure 7B). To determine if antibodies cleared parasite and prevented parasite antigens being presented to B cells, we took advantage of the fact that sporozoites rapidly infect the liver (<2 hours even after intradermal immunization), and are rapidly taken up into antigen presentation pathways in the spleen or draining lymph nodes ^39, 40^, this should therefore be circumvented if the antibody was delivered late. However, transfer of the antibody 4 hours after immunization only permitted a minimal expansion of memory B cells (Figure 7B). Therefore, these two general mechanisms cannot explain the observed negative antibody feedback. In contrast mAbs targeting either the N-terminal (5D5)^41^ or C-terminal (mAb15)^20^ domains of PfCSP had little or no inhibitory effect on responses by memory Igh^g2A10^ cells (Figure 7B) which is consistent with epitope masking being the mechanism of antibody feedback. This was true even when the 5D5 and mAb15 antibodies were added in significant excess (Figure 7C). Importantly, dissociation constants for the binding of 5D5, mAb15 and 2A10 to PfCSP are all in the same range (Supplementary Figure 6).

If epitope masking was the mechanism of antibody feedback in humans that received PfSPZ vaccine, then it should still be possible for responses to subdominant epitopes targeting other regions of PfCSP to continue to expand even if the immunodominant anti-repeat response has plateaued. We therefore separated responses in our vaccine cohort by specificity for the (NANP)_n_ repeat which is immune-dominant or the C terminus. As expected, responses to the (NANP)_n_ repeat were similar to those of whole PfCSP, plateauing after V2. Of note, however C terminal responses continued to increase after V3, suggesting antibody feedback was not acting on C-terminal responses (Figure 7D). In agreement with this there was correlation between V2 anti-(NANP)_n_ repeat titer and boost at V3 that was even stronger than that observed for total CSP responses (r^2^ = 0.74, p <0.0001; Figure 7E). In contrast no relationship was observed between antibody titer and the magnitude of boosting for C-terminal antibodies at any timepoint (Figure 7F). It is notable that C-terminal antibodies have been associated with protection by the RTS, S vaccine ^42, 43^. Overall these data strongly support the hypothesis that epitope masking rather than parasite clearance of Fc mediated immune regulator mechanisms account for antibody feedback not only in the mouse model but also in vaccinated individuals.

## Discussion

The standard approach for generating high titers of antibody has been a series of immunizations followed by periodic booster injections depending on the infection. For prevention of malaria it has proven difficult to achieve the necessary high titers to achieve high-level (>50%) protective antibody even with additional boosting. Using data from a human clinical trial combined with novel Ig-knockin mouse model we show that boosting is limited by antibody feedback which prevents antibodies to a single epitope reaching protective levels. While antibody feedback is a well-established immunological phenomenon ^44, 45, 46^, its role in limiting vaccine responses to complex pathogens has not been explored. Collectively our data provide support for vaccine approaches based on multiple protective epitopes delivered either sequentially or in parallel.

While we have principally used whole parasite vaccination with PfSPZ Vaccine (in humans) or Pb-PfCSP SPZ in mice for modelling antibody and B cell responses, the data herein may also explain some of the features of such responses to subunit vaccines for malaria. Notably in strong agreement with our data it has previously been reported that the subunit RTS,S vaccine induces anti-repeat responses that plateau after 2 immunizations, but responses to the Hepatitis B S antigen core of the subunit continue to rise upon a third dose ^6, 7^. Our data also show that boosting principally induces a short-lived PB response, which may temporarily allow the serum titers of anti-(NANP)_n_ antibodies to achieve protective levels; however, only a small fraction of responding cells differentiate to become long-lived BMPCs which may explain the short-lived protection by RTS,S even when additional late booster responses are given ^4, 5^. Understanding how to induce a larger number of long-lived BMPCs will thus be a critical challenge for future vaccination approaches. Interestingly we found that delayed boosting resulted in secondary GCs and higher antibody responses, which is consistent with findings that delayed administration of the RTS,S vaccine results in better maturation of the B cell response and protection than a series of boosts administered close together ^47^.

Our data may have implications for vaccines targeting pathogens other than malaria. It has been shown that responses to seasonal Influenza vaccination are inversely proportional to pre-existing anti-Influenza titers suggesting a role for antibody feedback in limiting responses ^48^. The seasonal Influenza vaccine is a trivalent vaccine containing the HA molecule from three circulating strains. When a novel trimeric cocktail is used for immunization that differs in some components from the previous vaccine, responses are more robust to the divergent antigens, consistent with a role for antibody feedback driving the diversification of the immune response^49^. For HIV, passive transfer studies in humans and vaccination studies in non-human primates suggest that concentrations of broadly neutralising antibodies (bnAbs) will be required to confer protection ^3, 50, 51, 52^. An additional problem is that bnAbs are highly mutated and hard to elicit by vaccination, current vaccine approaches therefore focus on giving a series of antigenically distinct Env trimers designed to stimulate germline precursors of bnAb producing plasma cells and focus the somatic hypermutation process ^53^. Antibody feedback may act as a double edge sword, simultaneously limiting overall responses but stimulating the diversification of the response at each step.

One striking finding is the mismatch between the amount of antibody required for protection and the amount required for feedback. In naïve mice, feedback inhibition was observed at serum concentrations of antibody of 1-10 μg/ml anti-PfCSP, which is equivalent to ∼7-70 × 10^−9^ M, which is a little above the previously reported K_d_ of 2A10 for PfCSP (2.7 × 10^−9^ M) ^13^. This would imply that antibodies can efficiently mask the presentation of their cognate epitopes to B cells at concentrations a little above their dissociation constant (K_d_). It is thus surprising that that serum concentrations of 2A10 of ∼200 μg/ml (>1 × 10^−6^ M) do not fully protect. This indicates that simple binding of antibody is insufficient for high-level protection. Thus, protective antibodies require some biological activity – either blocking the motility of sporozoite or blocking CSP function – to exert protection ^20^. One mechanism that may mitigate the low amount of antibody that inhibits B cells responses may be the induction of memory T cell help. Interestingly, in our experiments, antibody feedback appears less potent in an immune background compared to in naive mice. This indicates that there is help for memory B cells in immune mice that is not present in naïve mice, which is most likely attributable to T cells. Nonetheless the specificity and induction of follicular helper T cells by sporozoite vaccines has not been well-studied.

Our studies have been facilitated by the use of novel Ig-knockin mouse to dissect the B cell response to PfCSP. This tool permits adoptive transfer experiments and the tracking of memory B cells which would otherwise be challenging. The Igh^g2A10^ mouse is designed to carry B cells of endogenous affinity as it carries a germline-reverted IgH V_H_DJ_H_ rearrangement, which is free to pair with any light chain. Notably our estimate of the affinity of our cells for PfCSP (1.33 × 10^−7^ M) is more than 4 orders of magnitude lower than the affinity of MD4 mice (∼5×10^−12^ M) for their cognate antigen HEL determined by others using a similar approach^54^. Our Igh^g2A10^-knockin mouse also undergoes class switching and affinity maturation in a manner that appears to mirror these processes in human B cells. Due to the physiological nature of this mouse model we anticipate it will be a useful tool in future studies of the B cell response to PfSPZ and different vaccine modalities.

While antibody feedback is a well-established immunological phenomenon, its role in regulating vaccine induced responses has not been clearly dissected. Antibody feedback will probably be a critical challenge to the development of any vaccine where sustained high titers of neutralising antibody are required for protection. Collectively our data explain some of the challenges facing future vaccine development, but also offer some insights into how these challenges may be overcome.

## Materials and Methods

### Study subjects and clinical specimens

VRC 314 clinical trial (https://clinicaltrials.gov/; NCT02015091) ^10, 11^ was an an open-label evaluation of the safety, tolerability, immunogenicity and protective efficacy of Sanaria® PfSPZ Vaccine. Subjects, recruited at the University of Maryland, Baltimore in the high dose cohort received a total of three doses of 9×10^5^ PfSPZ intravenously at week 0, 8 and 16. Blood was drawn at the time of each immunization, as well as 7 d and 14 d after each immunization. Plasma and PBMCs were isolated from all samples at these timepoints.

### Isolation of plasmablasts

PBMCs isolated from blood samples collected 7 d after immunization with PfSPZ Vaccine were used fresh or frozen and thawed prior to staining for viability with Aqua LIVE/DEAD dye (Invitrogen) followed by surface staining and FACs sorting (full details of antibodies are given in Supplementary Table 1). PBs were gated according to Supplementary Figure 1A and sorted as single cells into 96-well PCR plates containing 20 µl/well of reverse transcriptase reaction buffer that included 5 µl of 5× first-strand cDNA buffer, 0.5 µl of RNAseOut (Invitrogen), 1.25 µl of dithiothreitol (DTT), 0.0625 µl of igepal and 13.25 µl of distilled H_2_O (Invitrogen) as previously described^21^.

### Production of recombinant immunoglobulins

Immunoglobulin-encoding genes of PBs were amplified through RT and nested PCR without cloning from RNA of single sorted cells as previously described^19, 21^. The amplified rearranged gene segments encoding variable regions were assembled into the corresponding linear full-length immunoglobulin heavy- and light-chain gene expression cassettes through PCR as previously described^19, 21^. Heavy and light chain linear cassettes were co-transfected in 293T cells using Effectene with enhancer (Qiagen)^19, 21^. Transfected cultures were incubated at 37 °C 5% CO_2_ for 3 d. Supernatants were harvested, concentrated and purified using HiTrap Protein A prepacked high-performance plates (GE Healthcare) for 20 min at room temperature on a shaker. Following wash with PBS and NaCl, eluates were neutralized with Trizma hydrochloride and buffer exchanged with PBS before determining antibody concentration using Nanodrop. Immunogenetic information was assigned to antibody sequences using Cloanalyst^55^ based on the IMGT immunoglobulin gene segment libraries (http://www.imgt.org)^56^. Sequences were deemed to be clonally related using Cloanalyst^55^ based on their V and J gene usage and CDR3 similarity.

### Screening of recombinant antibodies via ELISA

Recombinant monoclonal antibodies were screened for PfCSP reactivity using either ELISA or electrochemiluminescence via the mesoscale discovery (MSD) platform. For ELISA MaxiSorp ELISA plates (Thermo Scientific Nunc) were coated with 100 µl of rPfCSP (1 µg/ml) per well for 1 h at room temperature according to the manufacturer’s instructions (KPL). Coated plates were blocked with 100 µl of 1× blocking solution for 1 h at room temperature, which was followed by incubation with 100 µl of PfCSP monoclonal antibodies, mock transfection filtrate or control antibodies (VRC 01, a human anti-HIV-1 IgG1 as an isotype-matched negative control^57^; 2A10, a mouse monoclonal antibody specific for the (NANP)_n_-repeat region of PfCSP^24, 58^) at varying concentrations (0.00006–5.0 µg/ml). After 1 hr, plates were incubated with 100 µl/well of 1.0 µg/ml peroxidase-labeled goat anti–human IgG antibody (KPL). Plates were washed six times with PBS-Tween between each step. After a final wash, samples were incubated for about 15 min with the ABTS peroxidase (KPL) or Ultra TMB ELISA (Invitrogen) substrate. The optical density was read at 405 or 450 nm after addition of stopping solution (100 µl/well).

### Mesoscale Discovery (MSD) ELISA for PfCSP sera titers and screening of recombinant monoclonal antibodies

Streptavidin MSD 384 well plates (MSD) were first blocked with PBS + 5%BSA for 30min, washed five times, then coated with biotinylated antigen PfCSP-biotin, NANP Repeat-Biotin, NTerm-Biotin, or CTerm-Biotin) at 1 µg/mL in PBS + 1% BSA. After 1 hr, plates were washed and 10 µl of serially diluted sera (starting at 1:10, then by 5 fold dilutions) in PBS + 0.05% Tween-20 + 1%BSA, was added and incubated for 1hr. Alternatively, for recombinant antibody screening antibody concentrations were normalized to 1 µg/ml in PBS + 0.05% Tween-20 + 1% BSA prior to loading onto plates. After washing, plates were incubated for 1 hr with sulfo-tag goat anti-human IgG detection antibody (MSD) at 1ug/mL diluted in PBS, 0.05%Tween, 1%BSA. Plates were washed, and 1x Read T buffer (MSD) diluted in distilled water was added before analyzing with MSD SECTOR Imager 2400. The log of mean fluorescence intensity (MFI) is reported. All incubations were done at room temperature and all wash steps were performed 5 times.

### Mice; generation of Igh^g2A10^ knockin animals

C57BL/6NCrl were purchased from the Australian Phenomics Facility (Canberra, ACT, Australia). MD4 mice (C57BL/6-Tg(IghelMD4)4Ccg; MGI:2384162) and Blimp1^GFP/+^ (C57BL/6(Prdm1^tm1Nutt^; MGI: 3510704) mice were a kind gift from Carola Vinuesa (The Australian National University). FLPe deleter mice (B6.Cg-Tg(ACTFLPe)9205Dym/J; MGI: 3714491)^59^ were imported from Jackson laboratories (Bar Harbor, ME; stock number 005703). Igh^g2A10^ were generated by Ozgene Pty Ltd (Bentley, WA, Australia) via embryonic cell transformation. Briefly the predicted germline precursor gene of the heavy chain of the 2A10 antibody was synthesised and inserted into a plasmid carrying flanking regions corresponding to positions chr12:113430554 to chr12:113435542 (for the 5’homology arm) and chr12:113425551 to chr12:113428513 (for the 3’ homology arm) for targeting into the IgM locus. Upstream of the g2A10 gene was an Igh promoter and a neomycin cassette flanked by Frt sites for subsequent excision. Subsequently mice were crossed to FLPe deleter mice which constitutively express FLPe under the control of the actin promoter to generate mice Igh^g2A10^ mice lacking the Neomycin cassette. Mice were bred and maintained under specific pathogen free conditions in individually ventilated cages at the Australian National University. All animal procedures were approved by the Animal Experimentation Ethics Committee of the Australian National University (Protocol numbers: A2013/12 and A2016/17). All mice were 5-8 weeks old at the commencement of experiments. Within each experiment, mice were both age matched. Female mice were used throughout the experiments.

### Immunizations and Antibody Transfer

Mice were immunized IV with 5 × 10^4^ Pb-PfCSP SPZ crossed to an mCherry background to facilitate the identification of infected mosquitoes^27, 60^. Sporozoites were dissected by hand from the salivary glands of *Anopheles stephensi* mosquitoes and were irradiated (200kRad) using a MultiRad 225 (Flaxitron) irradiator prior to injection. For PfCSP immunization, 30µg rPfCSP^13^ in PBS was absorbed with Imject Alum (ThermoFisher) in a 2:1 ratio of antigen: adjuvant according to the manufacturer’s instructions and injected IP in a final volume of 150 µl.

Antibodies for passive transfer were injected IV at the stated doses. 5D5^41^ (mouse IgG1) was a kind gift of Gabriel Gutierrez (Leidos). CIS43^20^, VRC01^57^ and mAb15^20^ (all human IgG1), were expressed in-house at the Vaccine Research Center from Expi293T cells. 2A10^24, 58^ (mouse IgG2a) was prepared from hybridoma cell supernatants (Genscript). 2A10 LALA-PG antibodies carrying L234A, L235A and P329G substitutions in the IgG2A heavy chain^38^ were expressed from Expi293F cells (Genscript). LTF-2 (mouse IgG2b) was purchased from BioXCell.

### Kinetic binding assay using biolayer interferometry

Antibody binding kinetics were performed using biolayer interferometry on an Octet Red384 instrument (fortéBio) using streptavidin-capture biosensors (fortéBio) as previously described ^20^. PfCSP monoclonal antibody solutions were plated in solid black tilt-well 96-well plates (Geiger Bio-One). Loading of biotinylated rPfCSP was performed for 300 s, followed by dipping the biosensors into buffer (PBS + 1% BSA) for 60 s to assess baseline assay drift. Association with whole IgG (serially diluted from 33 to 0.5208 µM) was done for 300 s, followed by a dissociation step in buffer for 600 s. Background subtraction of nonspecific binding was performed through measurement of association in buffer alone. Data analysis and curve fitting were performed using Octet software, version 7.0. Experimental data were fitted with the binding equations describing a 1:1 heterologous ligand interaction. Global analyses of the complete data sets, assuming binding was reversible (full dissociation), were carried out using nonlinear least-squares fitting allowing a single set of binding parameters to be obtained simultaneously for all concentrations of a given monoclonal antibody dilution series.

### Flow Cytometry and sorting

Lymphocytes were isolated from the spleen and bone marrow of mice and were prepared into single cell suspensions for flow cytometric analysis and sorting. Bone marrow cells were flushed from femurs and tibias with FACs buffer in 27g syringes, whilst splenocytes were isolated by mashing spleens over 70 μm micron mesh filters. Red blood cells were lysed from cell suspensions with ACK lysis buffer (Sigma) and cells were washed twice with FACs wash prior to antibody staining. Cells were quantified during flow cytometry by the addition of CountBright Absolute Counting beads (Invitrogen) to sample suspensions. Details of antibodies are given in Supplementary Table 1, and details of generic gating strategies for mouse experiments are given in Supplementary Figure 2. (NANP)_9_ tetramers were prepared in house by mixing biotinylated (NANP)_9_ peptide with streptavidin conjugated PE or APC (Invitrogen) in the a 4:1 molar ratio. Flow-cytometric data was collected on a BD Fortessa or X20 flow cytometer (Becton Dickinson) and analyzed using FlowJo software (FlowJo). A BD FACs Aria I or II (Becton Dickinson) machine was used for FACS sorting of cells.

### Cell purification and adoptive transfers

For primary immunizations the number of naïve tetramer^+^ cells were quantified from Igh^g2A10^ splenocytes via flow cytometry. The concentration of donor splenocytes were then adjusted to deliver 1-2 × 10^4^ tetramer^+^ Igh^g2A10^ CD19+ B cells to each recipient mouse in 100 μl via IV injection. Mice were immunized 1-2 days after adoptive transfer of naïve splenocytes. Memory cells for adoptive transfer experiments were enriched from mice that had received naïve Igh^g2A10^ B cells and been immunized with either Pb-PfCSP SPZ or PfCSP. 8-10 weeks after immunization, memory cell donor mice were culled and single cell preparations of lymphocytes from the spleen were made. Splenocytes were treated with FC-block (anti-CD16/32, Biolegend) and then incubated with lineage specific biotin conjugates (anti-GL7, anti-CD138, anti-CD4, and anti-CD8; see supplementary table 1 for details of clones and suppliers) to remove T cells, PBs and germinal centre B cells. Afterwards, splenocytes were incubated with anti-biotin microbeads (Miltenyi), and then put through a magnetic LS column (Miltenyi), according to the manufacturer’s directions. A cell suspension enriched in Igh^g2A10^ memory B cells was collected in the runoff after passing through the column (Supplementary Figure 5). A fraction of the sample was stained for flow cytometric analysis to quantify the number of CD45.1^+^Tet^+^ memory cells, and to confirm that depletion of other B cell subsets and helper T cells had worked. The concentration of cells was adjusted such that 5 × 10^2^ Igh^g2A10^ memory cells were then transferred to recipient mice IV in a volume of 100 μl.

### ELISA for the detection of anti-PfCSP antibodies in mice

Concentrations of PfCSP specific antibodies in the sera of mice after immunization were measured using solid phase ELISA. Briefly, Nunc Maxisorp Plates (Nunc-Nucleon) were coated overnight with 1ug/ml streptavidin followed by binding of biotinylated (NANP)_9_ peptide for 1 hour. After blocking with 1% BSA, serial dilutions of the antibodies were incubated on the plates for 1 hour and after washing, incubated with HRP conjugated anti-mouse IgG or anti-mouse IgM antibodies (KPL). Plates were developed for 15-20 minutes with ABTS 2-Component Peroxidase Substrate Kit (KPL), and read at 405nm using a Tecan Infinite 200Pro plate reader. IgM responses were analysed as the area under the absorbance curve (AUC). IgG concentrations were calculated from a standard curve that was generated from serial dilutions of 2A10 (starting at 1μg/ml).

To assess circulating levels of passively transferred PfCSP-specific monoclonal antibodies, mice were bled via the tail plexus immediately before challenge with infectious mosquito bites. ELISA was performed on mouse serum as previously described using plates coated as above. A standard curve for each monoclonal antibody was generated using eight 3-fold dilutions of monoclonal antibody starting at 1 μg/ml. Serum samples were applied at a series of dilutions from 1:500-1:4500 in blocking buffer. The concentrations were commonly calculated off the values from the ∼1:1500 dilution. However, if the sample was uncommonly low/high then other dilutions were used for the calculation (providing they sat within the exponential range of the monoclonal standard curve).

### ELISpot for the detection of PfCSP specific plasmablasts

Sterile MultiScreen*™*-HA plates (Millipore) were coated with PfCSP at 2.5 μg/ml in PBS, and left overnight at 4C. After washing with sterile PBS, plates were blocked with complete RPMI 1640 (10% FCS, 2 mM L-glutamine, 1 mM Na-Pyruvate, 100 U/ml Penicillin/Streptomicin, 5mM HEPEs, 20 μg/ml Gentamicin and 50 μM *β*-mercaptoethanol) for 3 hours at 37C. Sorted bone marrow derived plasma cells were added to selected wells, and incubated overnight at 37C. The following day plates were washed with wash buffer, and incubated for 3 hours with HRP conjugated anti-mouse IgG (KPL). After washing with wash buffer, plates were developed using stable DAB (Invitrogen) for 20 minutes. The number of spots was counted manually in each well by two individuals blinded to the experimental groups.

### In vivo protection in C57BL/6 mice with chimeric Pb-PfCSP SPZ

For the mosquito bite challenge, female *A. stephensi* mosquitoes were allowed to feed on 8-week-old C57BL/6 mice infected with blood-stage Pb-PfCSP-mCherry. 21 days after feeding mosquitoes were chilled on ice and sorted for infection. The abdomens and thoraxes of infected mosquitoes glow red under green fluorescent light due to the presence of mCherry^+^ midgut and salivary gland sporozoites facilitating sorting. Mice were challenged with ∼5 infected mosquitoes per mouse. C57BL/6 mice were injected IV with monoclonal antibodies as stated in the relevant figure legends. Ten minutes later, mice were anesthetized with Ketamine-Xylazine (100 mg/kg and 10 mg/kg respectively), and the infected mosquitoes were allowed to feed on mice for ∼30 min, after which mosquito abdomens were visually inspected for blood, indicating the mosquito has bitten. Mouse parasitemia was assessed daily through flow cytometry from day 4 through day 14 post-infection. A mouse was considered patent once parasitemia reached >0.1%.

### Single cell RNA-seq to sequence recombined Ig V(D)J chains from mice

Single Cell RNA sequencing was performed using a SMARTseq 2 protocol^33^ with the following modifications. Cells were sorted into plates with wells containing 1ul of the cell lysis buffer, 0.5 μl dNTP mix (10 mM) and 0.5 μl of the oligo-dT primer at 5 μM. We then reduced the amount reagent used in the following reverse-transcription and PCR amplification step by half. The concentration of the IS PCR primer was also further reduced to 50 nM. Due to the low transcriptional activity of memory B cells, we increased the number of PCR cycles to 28. Sequencing libraries were then prepared using the Nextera XT Library Preparation Kit with the protocol modified by reducing the original volumes of all reagents in the kit by 1/5^th^. Sequencing was performed on the Illumina NextSeq sequencing platform.

To determine the antigen-specific BCR repertoire, we made use of VDJpuzzle ^61^ to reconstruct full-length heavy and light chains from each cell. From this we were able to determine V region usage and mutation frequency.

### Experimental design and statistical analysis

Details of specific statistical tests and experimental design are given in the relevant figure legends. Mouse experiments had 3-5 mice per group and were performed either in duplicate or triplicate. All data points are plotted from all replicate experiments, though for one human subject (figure 1B/7B)) data were excluded from subsequent statistical analysis as these individuals did not respond to immunization; excluded data points are marked by #. In most instances analysis was performed in R (The R Foundation for Statistical Computing) on the pooled data from all replicate experiments. Where data was pooled from multiple experiments, each experiment was included as a blocking factor in the analysis. Where data are plotted on a log-scale data were log-transformed prior to analysis. For certain experiments and the analysis of some human data blocking factors did not need to be accounted for and analysis was performed in GraphPad Prism 7. Abbreviations for p values are as follows: p < 0.05 = *, p < 0.01 = **, p < 0.001 = ***, p < 0.0001 = ****; with only significant p values shown. With the exception of ELISpot counting, no blinding or randomization was performed, however all other readouts (ELISA, flow cytometry and sequencing) are objective readouts that are not subject to experimental bias.

## Author Contributions

Conceptualization: H.A.M., A.H.I., R.A.S., I.A.C.; Investigation: H.A.M., A.H.I., H.J.S., B.J.F., Y.C., D.C., K.L., S.C., N.K., B.K.L.S., M.B., I.A.C.; Formal analysis: H.A.M., A.H.I., H.J.S., K.W., M.B., I.A.C.; Resources: S.L.H.; Writing-original draft preparation: H.A.M. and I.A.C.; Writing - review and editing: A.H.I., M.B. and R.A.S.. Project administration R.A.S. and I.A.C.; Funding acquisition: S.L.H., R.A.S. and I.A.C.

## Acknowledgements

This work was supported by start-up funds from the Australian National University to I.A.C. and NHMRC project grant support to I.A.C. (GNT1158404). Production and characterization of PfSPZ Vaccine were supported in part by National Institute of Allergy and Infectious Diseases Small Business Innovation Research Grants 5R44AI055229-11 (to S.L.H.), 5R44AI058499-08 (to S.L.H.), and 5R44AI058375-08 (to S.L.H.). We would like to thank the University of Maryland study volunteers from malaria clinical trial VRC314. We would like to that Harpreet Vohra and Michael Devoy of the Imaging and Cytometry Facility at the Australian National University for assistance with flow cytometry and sorting. We thank Morgan Gladden, Anthony Monroe, Ruijun Zhang, Minyue Wang and Joshua Beem of the Duke Human Vaccine Institute for assistance with gene amplification, antibody production and immunogenetics analysis. Special thanks to Joe R. Francica for technical support and guidance with antibody affinity measurements. We would like to acknowledge the support of the flow cytometry core at the Vaccine Research Center particularly David Ambrozak for assistance with PB sorting.

