## Supplementary material for "Antibody feedback limits the expansion of cognate memory B cells but drives the diversification of vaccine-induced antibody responses": Supplmentary Information

**Supplementary Information**

This supplementary information contains:

Supplementary Table 1: Antibodies used in this study

Supplementary Dataset 1: Sequence data from analysis of plasmablasts from PfSPZ vaccinated volunteers

Supplementary Figures 1-6

**Supplementary Table 1: Antibodies used in this study**

**A. Anti-mouse antibodies for flow cytometry and magnetic column sorting**

| Antigen | Conjugate | Clone | Source | Catalogue | Concentration | Dilution |
|---|---|---|---|---|---|---|
| Anti-mouse B220 | APC | RA3-6B2 | Biolegend | 103212 | 0.2 mg/mL | 1/400 |
| Anti-mouse B220 | BV605 | RA3-6B2 | BioLegend | 103244 | 0.2 mg/mL | 1/400 |
| Anti-mouse B220 | PE-Cy7 | RA3-6B2 | BD Pharmigen | 552772 | 0.2mg/mL | 1/200 |
| Anti-mouse CD3 | PerCP Cy5.5 | 17A2 | Biolegend | 100218 | 0.2 mg/mL | 1/200 |
| Anti-mouse CD4 | APC | RM4-5 | Biolegend | 100516 | 0.2 mg/mL | 1/200 |
| Anti-mouse CD4 | Biotin | GK1.5 | Biolegend | 100404 | 0.5 mg/mL | 1/200 |
| Anti-mouse CD5 | APC | 537.3 | eBioscience | 17-0051-82 | 0.2mg/mL | 1/400 |
| Anti-mouse CD8a | Biotin | 53-6.7 | Biolegend | 100704 | 0.5 mg/mL | 1/200 |
| Anti-mouse CD8a | BV421 | 53-6.7 | Biolegend | 100738 | 0.2 mg/mL | 1/200 |
| Anti-mouse CD8a | FITC | 5H10-1 | Biolegend | 100804 | 0.5 mg/mL | 1/200 |
| Anti-mouse CD11a | PE Cy7 | M17/4 | Biolegend | 101122 | 0.2 mg/mL | 1/200 |
| Anti-mouse CD11b | FITC | M1/70 | Biolegend | 101206 | 0.5 mg/mL | 1/100 |
| Anti-mouse CD11b | PerCP Cy5.5 | M1/70 | Biolegend | 101228 | 0.2 mg/mL | 1/200 |
| Anti-mouse CD11c | PerCP Cy5.5 | N418 | Biolegend | 117328 | 0.2 mg/mL | 1/200 |
| Anti-mouse CD19 | BUV395 | 1D3 | BD Horizon | 563557 | 0.2 mg/mL | 1/200 |
| Anti-mouse CD21/35 | FITC | 7G6 | BD Pharmigen | 553818 | 0.5mg/ml | 1/400 |
| Anti-mouse CD23 | Pac Blue | B3B4 | Biolegend | 101616 | 0.5mg/ml | 1/200 |
| Anti-mouse CD24 | Pac Blue | M1/69 | Biolegend | 101820 | 0.5 mg/mL | 1/200 |
| Anti-mouse CD28 | APC | E18 | Biolegend | 122016 | 0.2mg/mL | 1/200 |
| Anti-mouse CD38 | A700 | 90 | eBioscience | 56-0381-82 | 0.2 mg/mL | 1/200 |
| Anti-mouse CD43 | BB515 | 57 | BD Horizon | 564646 | 0.2mg/mL | 1/200 |
| Anti-mouse CD44 | Pac Blue | IM7 | Biolegend | 103020 | 0.5mg/ml | 1/400 |
| Anti-mouse CD45.1 | BV510 | A20 | Biolegend | 110741 | 0.2mg/ml | 1/200 |

| | | | | | | |
|---|---|---|---|---|---|---|
| Anti-mouse CD45.1 | FITC | A20 | Biolegend | 110706 | 0.5 mg/mL | 1/200 |
| Anti-mouse CD45.1 | PE | A20 | Biolegend | 110708 | 0.2mg/mL | 1/200 |
| Anti-mouse CD45.2 | FITC | 104 | Invitrogen | 11-0454-82 | 0.5 mg/mL | 1/200 |
| Anti-mouse CD45.2 | PE | 104 | Biolegend | 109807 | 0.2 mg/mL | 1/200 |
| Anti-mouse CD80 | BV421 | 16-10A1 | Biolegend | 104725 | 0.2mg/mL | 1/400 |
| Anti-mouse CD80 | PE CY7 | 16-10A1 | Biolegend | 104734 | 0.2 mg/mL | 1/200 |
| Anti-mouse CD93 | APC | AA4.1 | eBioscience | 17-5892-82 | 0.2 mg/mL | 1/100 |
| Anti-mouse CD138 | Biotin | 281-2 | Biolegend | 142512 | 0.5 mg/mL | 1/200 |
| Anti-mouse CD138 | BV510 | 281-2 | Biolegend | 142521 | 0.2 mg/mL | 1/300 |
| Anti-mouse CD138 | PE Cy7 | 281-2 | Biolegend | 142514 | 0.2 mg/mL | 1/300 |
| Anti-mouse CD273 (PDL2) | APC | TY25 | Biolegend | 107210 | 0.2 mg/mL | 1/200 |
| Anti-mouse GL7 | A.488 | GL7 | Biolegend | 144612 | 0.5 mg/mL | 1/400 |
| Anti-mouse GL7 | Biotin | GL7 | Biolegend | 144616 | 0.5 mg/mL | 1/200 |
| Anti-mouse GL7 | eFluor450 | GL-7 | Invitrogen | 48-5902-82 | 0.2 mg/mL | 1/100 |
| Anti-mouse IgD | BV605 | 11-26c.2a | Biolegend | 405727 | 0.2 mg/mL | 1/400 |
| Anti-mouse IgM | APC-eFluor780 | II/41 | Invitrogen | 47-5790-82 | 0.2 mg/mL | 1/200 |
| Anti-mouse Ly-6G/Ly-6C (GR1) | PerCP Cy5.5 | RB6-8C5 | Biolegend | 108428 | 0.2 mg/mL | 1/200 |
| Anti-mouse NK1.1 | PE-CY7 | PK136 | Biolegend | 108714 | 0.2mg/mL | 1/200 |
| TruStain fcX (rat anti-mouse CD16/32): | - | 93 | Biolegend | 101320 | 0.5 mg/mL | 1/50 |
| 7AAD Cell Viability Dye | PerCP Cy5.5 | - | Biolegend | 420404 | 50 µg/mL | 1/100 |

**B. Anti-human antibodies for flow cytometry**

| Antigen | Conjugate | Clone | Source | Catalogue |
|---|---|---|---|---|
| Anti-human CD19 | FITC | HIB19 | BD Bioscience | 555412 |
| Anti-human CD20 | APC-Cy7 | L27 | BD Bioscience | 335794 |
| Anti-human CD27 | APC | O323 | ThermoFisher | 17-0279-42 |

| | | | | |
|---|---|---|---|---|
| Anti-human CD3 | PE-Cy7 | SK7 | BD Bioscience | 557851 |
| Anti-human CD38 | PE | HIT | BD Bioscience | 555460 |
| Live Dead Aqua | - | - | Invitrogen | L34966 |

**C. Antibodies for ELISA and ELIspot**

| Antigen | Conjugate | Source | Catalogue | Concentration | Dilution |
|---|---|---|---|---|---|
| Anti-human IgG (H+L) - produced in goat | HRP | Seracare | 5220-0330 | 1mg/ml | 1/2000 |
| Anti-mouse IgG (H+L) - produced in goat | HRP | Seracare | 5220-0341 | 1mg/ml | 1/2000 |
| Anti-mouse IgM (μ) - produced in goat | HRP | Seracare | 074-1803 | 1mg/ml | 1/2000 |

**Supplementary Dataset 1: Sequence data from analysis of plasmablasts from PfSPZ vaccinated volunteers.** Complete details of sequences and specificities for all plasmablasts isolated from vaccinated donors. CSP reactive cells are denoted in grey if the sole representative of a clone. Individual members of expanded clones are colour coded according to the clone they belong to.

#### Supplementary Figure Legends

**Supplementary Figure 1: Isolation and screening of human plasmablasts.**

PfSPZ vaccinated individuals were bled one week after each boost and plasmablasts sorted, their BCR genes were cloned and expressed and measured for reactivity with CSP by ELISA or electrochemiluminescence. A. Gating strategy for the single cell sorting of human plasmablasts. B. CSP reactivity of each antibody cloned from the plasmablasts at the different time. CSP reactivity was measured by ELISA (615 all timepoints; 606 V2) or electrochemiluminescence (608 all timepoints; 606 V1 and V3). For both techniques background cut-off for CSP reactivity was set at Median +4MAD. Data are expressed as % of maximum value with the background subtracted.

**Supplementary Figure 2: Generation and phenotyping of the Igh$^{g2A10}$ mouse.**

A. Schematic of the insertion of the rearranged heavy chain VDJ exon upstream of the IgM locus. B. Gating strategy for phenotyping of splenic B cell subsets (i) and relative percentages of splenic lymphocyte subsets in Igh$^{g2A10}$ mice compared with C57BL/6 mice (ii). C. Gating strategy for B cell subsets within bone marrow (i) and relative percentages (ii). D. Gating strategy for B cell subsets within peritoneal cavity (i) and relative percentages (ii). G. Strategy for identification of Igh$^{g2A10}$ B cells following adoptive transfer to congenic recipient mice. H. Strategy for identification of tetramer$^{+}$ Igh$^{g2A10}$ long lived plasma cells in the bone marrow of recipient mice when using congenic markers to sort cells (i) or using Igh$^{g2A10}$ *Blimp*$^{GFP/+}$ reporter cells (ii). All analyses between the Igh$^{g2A10}$ and C57BL/6 performed using a Student's t-test for each pairwise comparison; means ± s.d shown.

**Supplementary Figure 3: CSP-reactive Igh$^{g2A10}$ cells undergo affinity maturation.**

Single tetramer$^{+}$ Igh$^{2A10}$ BMPCs were sorted from mice 1 month after transfer of 2 x10$^{4}$ Igh$^{g2A10}$ cells and vaccination with 5 x 10$^{4}$ Pb-PfCSP SPZ and the BCR sequenced. The locations of mutations in the heavy chain (A) and light chain (B) are shown for IgG$^{+}$ and IgM$^{+}$ plasmablasts and compared to the sites of mutations in the 2A10 monoclonal antibody. Mutations that are annotated at the top of each are those that have been previously validated as increasing affinity over the germline unmutated precursor.

**Supplementary Figure 4: Isolation and phenotyping of PfCSP-specific memory B cells.**

A. Phenotype of memory B cells generated 2 months after transfer 2 x10$^{4}$ Igh$^{g2A10}$ cells and immunization with either 5 x 10$^{4}$ Pb-PfCSP SPZ or 30 μg PfCSP in Alum. Fluorescence-minus-one (FMO) staining controls for CD80 and PDL2 also shown. B. Relative percentage of PDL2$^{+}$ and CD80$^{+}$ memory cell subsets generated from either immunogen. C. Memory B cells prepared as in A. were depleted of T cells, GC B cells and Plasmablasts and analysed to confirm the efficiency of enrichment. Panels show the Igh$^{g2A10}$ B cells populations present in the immunised recipients with and without the depletion of non-memory B cell using magnetic columns. Once memory Igh$^{g2A10}$ cell suspensions were depleted of other contaminating B cell populations they were then used for adoptive transfer to naïve or immune recipients.

**Supplementary Figure 5: Delayed boosting allows enhanced recall responses.**

A. Schematic of the vaccination schedule for the experiment. B. Concentration of IgG anti-$(NANP)_9$ antibodies in the sera of mice immunized as in A; each icon denotes an individual serum sample with bars showing means ± sd. C. Representative flow cytometry plots for the identification of $Igh^{g2A10}$ B populations within the spleens of recipient mice. Summary data for the analysis of spleen PBs (D), spleen GC B cells (E), spleen memory B cells (F) and BMPCs (G) at the indicated timepoints; data are pooled from 2 replicate experiments, analysis was by one-way ANOVA including the experiment as a blocking factor, bars show means ± s.d..

**Supplementary Figure 6: Binding of repeat and non-repeat specific monoclonal antibodies to PfCSP by bilayer interferometry.**

Avidity of 2A10, 5D5 and mAb15 antibodies for rPfCSP. Antibody binding curves are shown in black (raw data). Data were fitted (dotted red lines) with the binding equations describing a 1:1 heterologous ligand interaction. Serial concentrations of antibodies used are as denoted to the right of each graph (n = 3, representative experiment is shown). Table shows the inferred binding kinetics of each antibody for PfCSP and associated errors.

Supplementary Figure 1

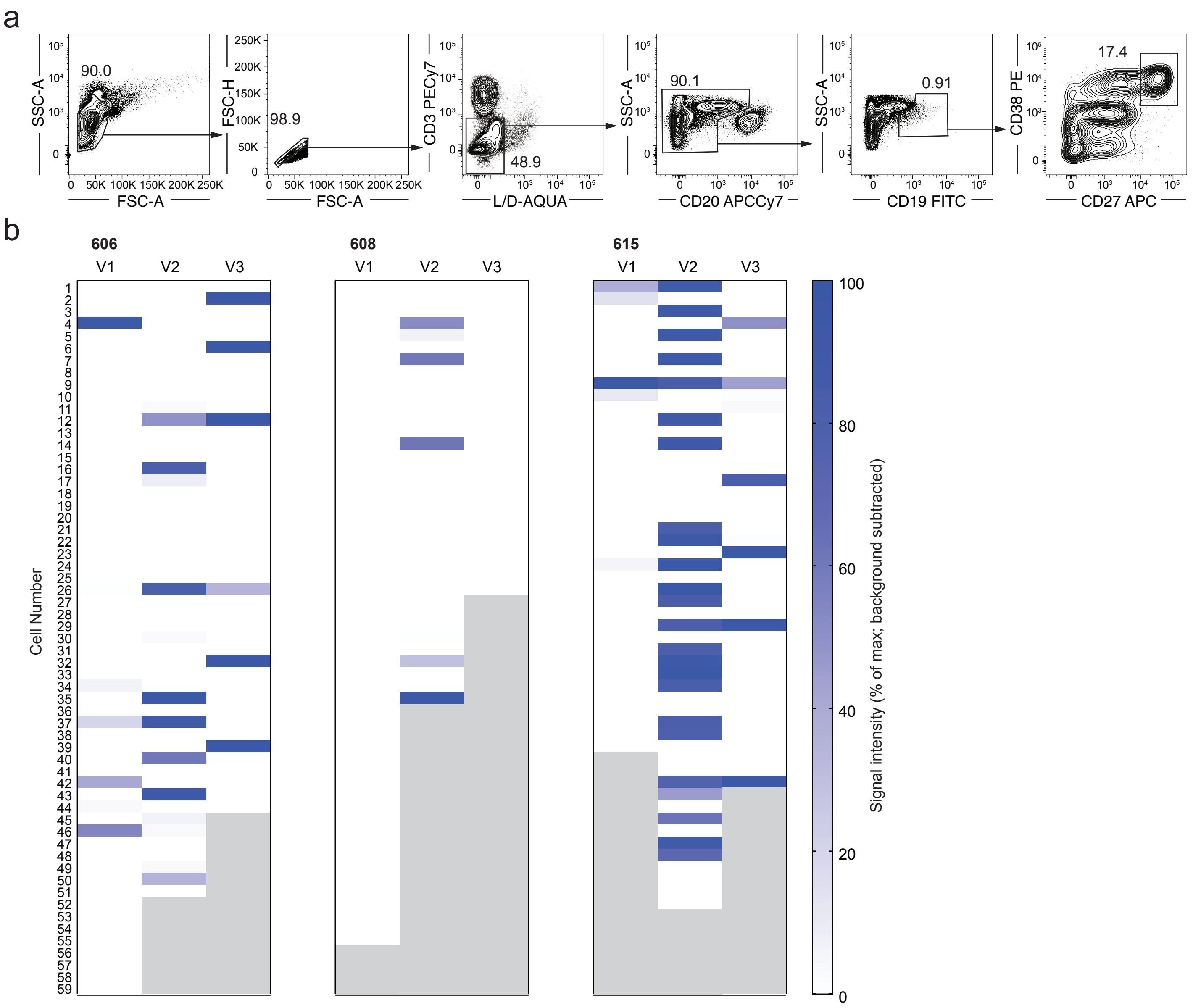


**Supplementary Figure 1: Isolation and screening of human plasmablasts.**
PfSPZ vaccinated individuals were bled one week after each boost and plasmablasts sorted, their BCR genes were cloned and expressed and measured for reactivity with CSP by ELISA or electrochemiluminescence. A. Gating strategy for the single cell sorting of human plasmablasts. B. CSP reactivity of each antibody cloned from the plasmablasts at the different time. CSP reactivity was measured by ELISA (615 all timepoints; 606 V2) or electrochemiluminescence (608 all timepoints; 606 V1 and V3). For both techniques background cut-off for CSP reactivity was set at Median +4MAD. Data are expressed as % of maximum value with the background subtracted.

### Supplementary Figure 2

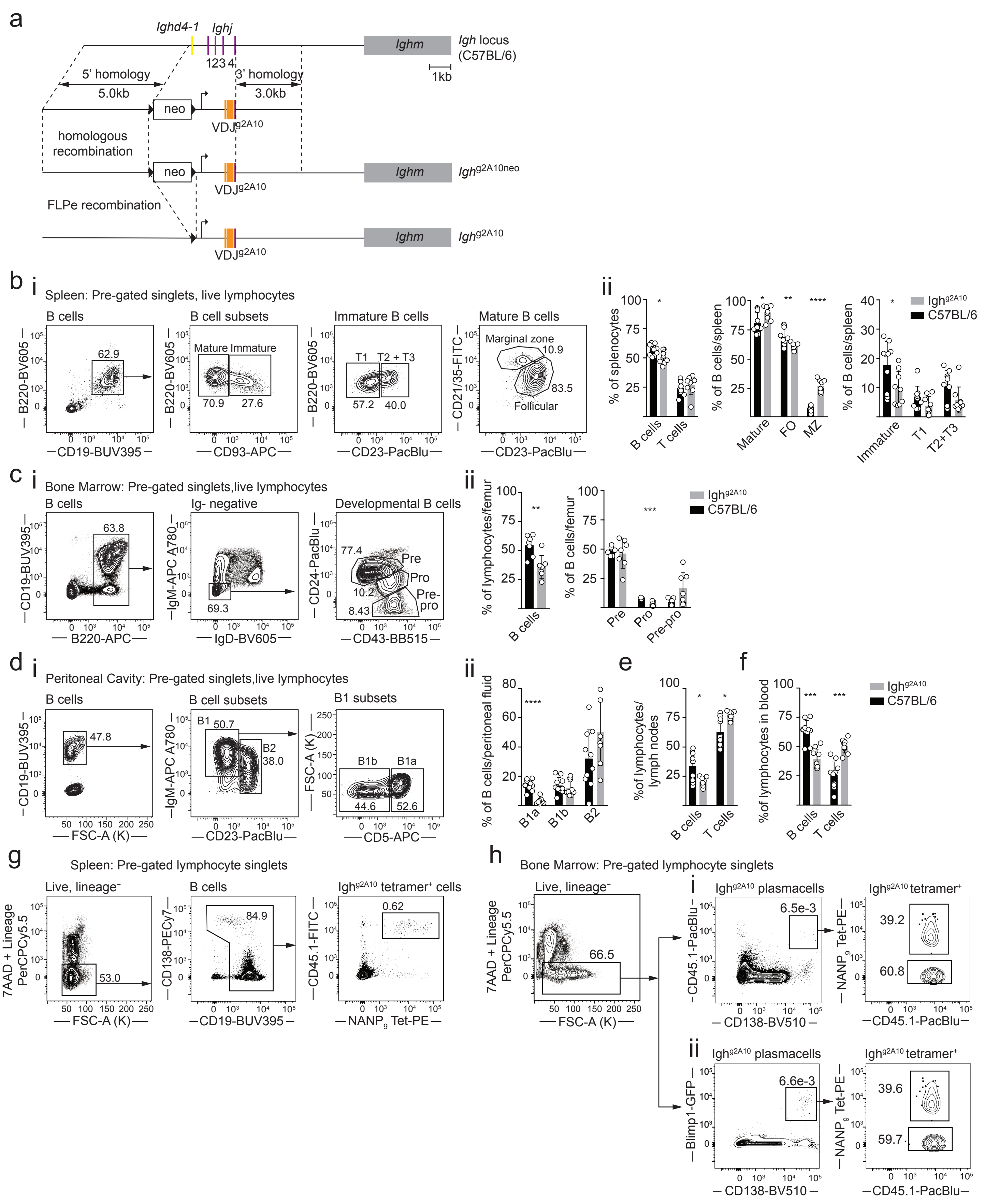

**Supplementary Figure 2: Generation and phenotyping of the Igh$^{g2A10}$ mouse.**

A. Schematic of the insertion of the rearranged heavy chain VDJ exon upstream of the IgM locus. B. Gating strategy for phenotyping of splenic B cell subsets (i) and relative percentages of splenic lymphocyte subsets in Igh$^{g2A10}$ mice compared with C57BL/6 mice (ii). C. Gating strategy for B cell subsets within bone marrow (i) and relative percentages (ii). D. Gating strategy for B cell subsets within peritoneal cavity (i) and relative percentages (ii). G. Strategy for identification of Igh$^{g2A10}$ B cells following adoptive transfer to congenic recipient mice. H. Strategy for identification of tetramer+ Igh$^{g2A10}$ long lived plasma cells in the bone marrow of recipient mice when using congenic markers to sort cells (i) or using Ighg2A10 BlimpGFP/+ reporter cells (ii). All analyses between the Igh$^{g2A10}$ and C57BL/6 performed using a Student's t-test for each pairwise comparison; means  s.d shown.

#### Supplementary Figure 3

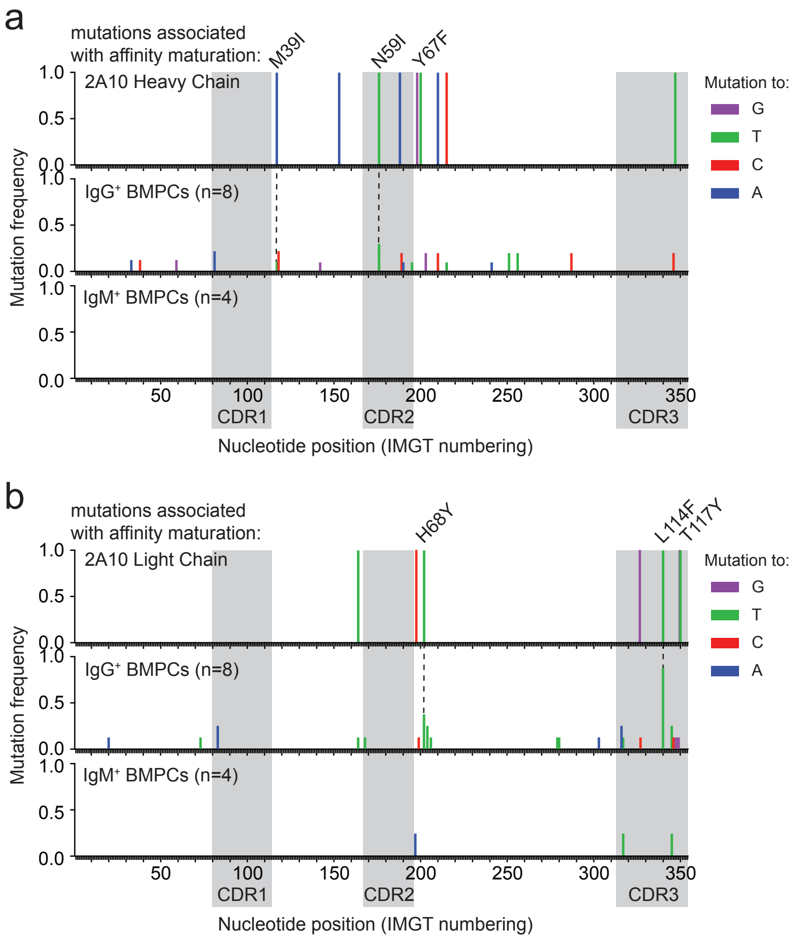


**Supplementary Figure 3: CSP-reactive Igh$^{g2A10}$ cells undergo affinity maturation.**
Single tetramer+ Ig$^{h2A10}$ BMPCs were sorted from mice 1 month after transfer of 2 x10$^{4}$ Igh$^{g2A10}$ cells and vaccination with 5 x 104 Pb-PfCSP SPZ and the BCR sequenced. The locations of mutations in the heavy chain (A) and light chain (B) are shown for IgG+ and IgM+ plasmablasts and compared to the sites of mutations in the 2A10 monoclonal antibody. Mutations that are annotated at the top of each are those that have been previously validated as increasing affinity over the germline unmutated precursor.

Supplementary Figure 4

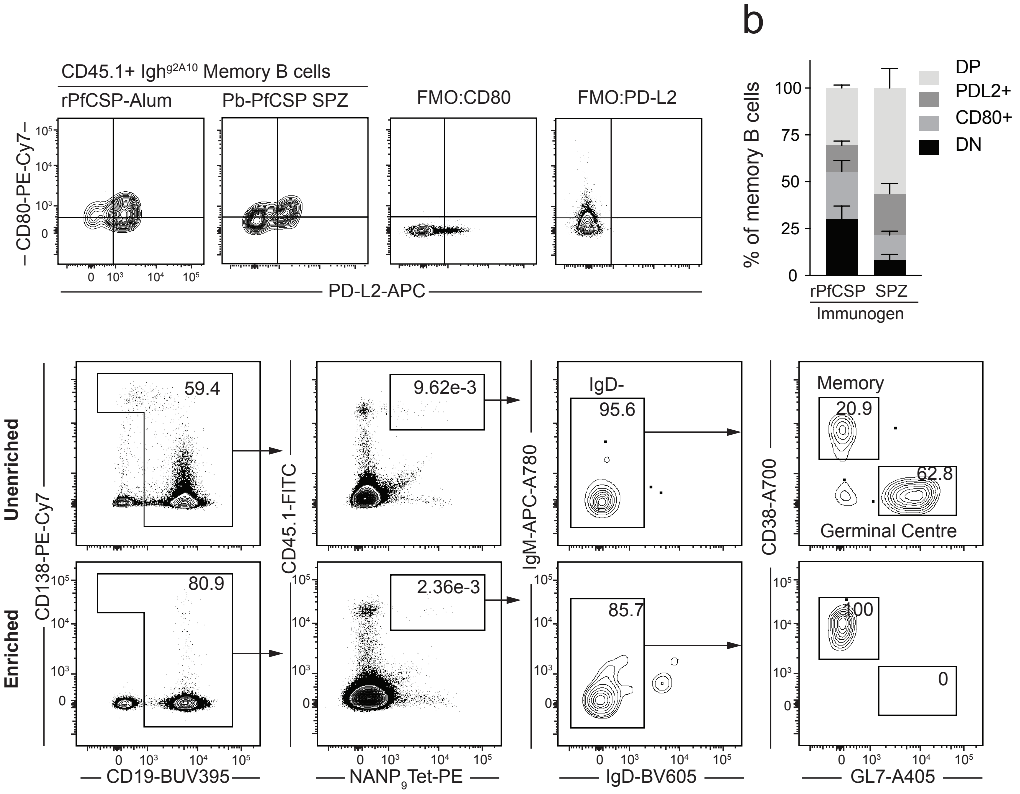


**Supplementary Figure 4: Isolation and phenotyping of PfCSP-specific memory B cells**
A. Phenotype of memory B cells generated 2 months after transfer 2 x104 Igh$^{g2A10}$ cells and immunization with either 5 x 104 Pb-PfCSP SPZ or 30 g PfCSP in Alum. Fluorescence-minus-one (FMO) staining controls for CD80 and PDL2 also shown. B. Relative percentage of PDL2+ and CD80+ memory cell subsets generated from either immunogen. C. Memory B cells prepared as in A. were depleted of T cells, GC B cells and Plasmablasts and analysed to confirm the efficiency of enrichment. Panels show the Igh$^{g2A10}$ B cells populations present in the immunised recipients with and without the depletion of non-memory B cell using magnetic columns. Once memory Igh$^{g2A10}$ cell suspensions were depleted of other contaminating B cell populations they were then used for adoptive transfer to naïve or immune recipients.

Supplementary Figure 5

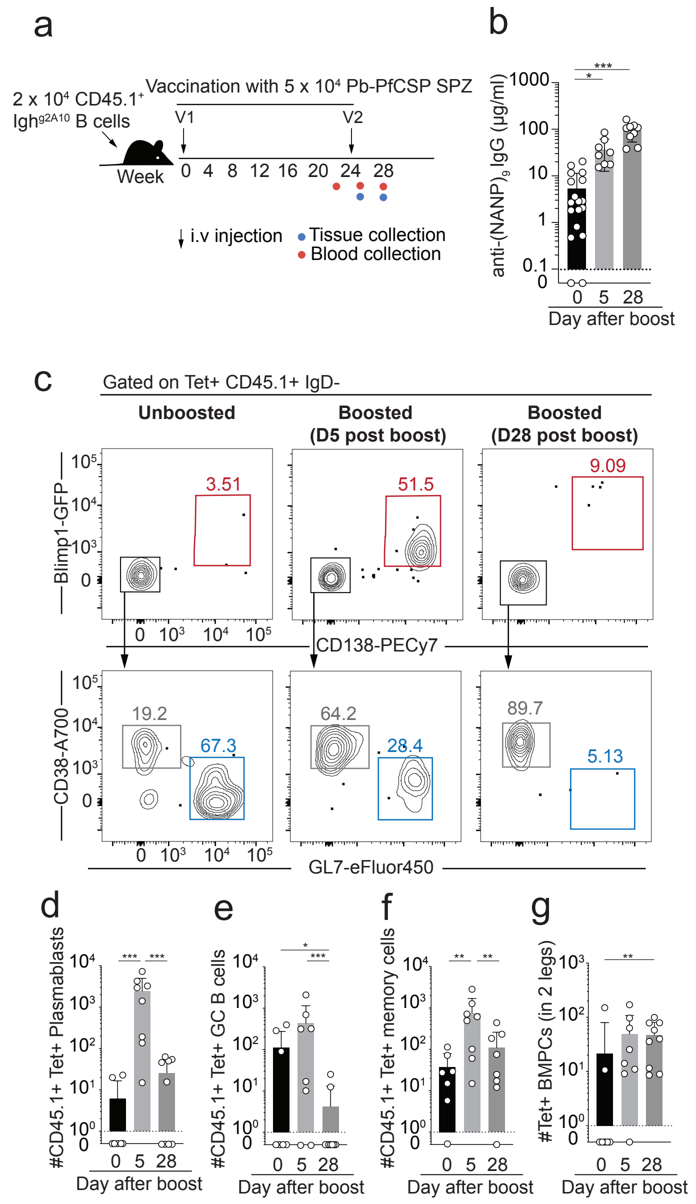


**Supplementary Figure 5: Delayed boosting allows enhanced recall responses.**
A. Schematic of the vaccination schedule for the experiment. B. Concentration of IgG anti-(NANP)$_9$ antibodies in the sera of mice immunized as in A; each icon denotes an individual serum sample with bars showing means  sd. C. Representative flow cytometry plots for the identification of Igh$^{g2A10}$ B populations within the spleens of recipient mice. Summary data for the analysis of spleen PBs (D), spleen GC B cells (E), spleen memory B cells (F) and BMPCs (G) at the indicated timepoints; data are pooled from 2 replicate experiments, analysis was by one-way ANOVA including the experiment as a blocking factor, bars show means  s.d..

Supplementary Figure 6

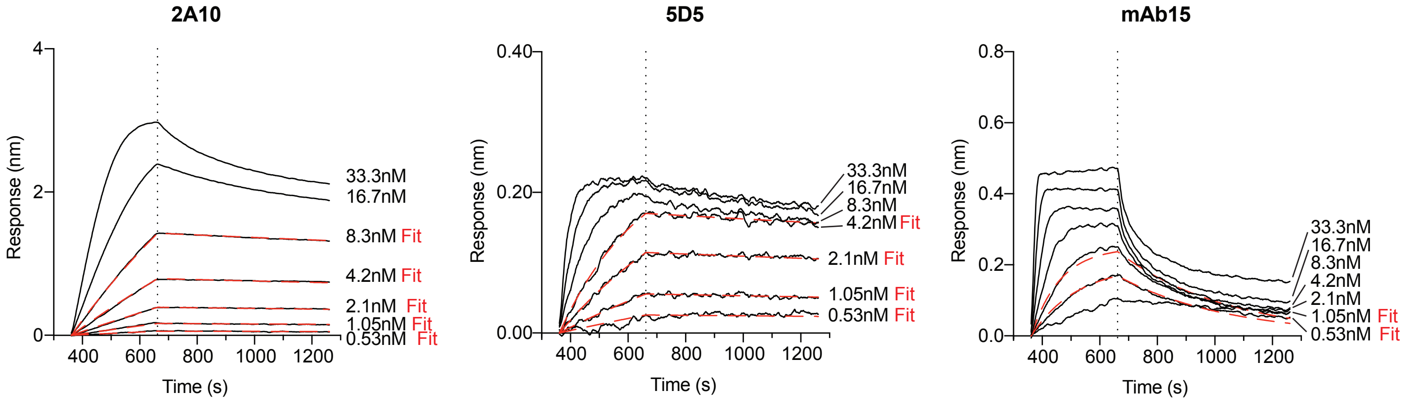


| mAb ID | $K_D$ (M) | $K_D$ Error | $k_{on}$(1/Ms) | $k_{on}$ Error | $k_{dis}$(1/s) | $k_{dis}$ Error |
|---|---|---|---|---|---|---|
| **2A10** | $1.05 \times 10^{-9}$ | $4.08 \times 10^{-11}$ | $1.20 \times 10^{5}$ | $3.98 \times 10^{3}$ | $1.27 \times 10^{4}$ | $2.55 \times 10^{6}$ |
| **5D5** | $1.76 \times 10^{-10}$ | $1.43 \times 10^{-11}$ | $7.91 \times 10^{5}$ | $3.36 \times 10^{4}$ | $1.39 \times 10^{4}$ | $9.67 \times 10^{6}$ |
| **mAb15** | $7.42 \times 10^{-10}$ | $4.47 \times 10^{-11}$ | $3.54 \times 10^{6}$ | $2.06 \times 10^{5}$ | $2.62 \times 10^{3}$ | $4.18 \times 10^{5}$ |

**Supplementary Figure 6: Binding of repeat and non-repeat specific monoclonal antibodies to PfCSP by bilayer interferometry**
Avidity of 2A10, 5D5 and mAb15 antibodies for rPfCSP. Antibody binding curves are shown in black (raw data). Data were fitted (dotted red lines) with the binding equations describing a 1:1 heterologous ligand interaction. Serial concentrations of antibodies used are as denoted to the right of each graph (n = 3, representative experiment is shown). Table shows the inferred binding kinetics of each antibody for PfCSP and associated errors.
